# Rax1/2 promote memory of growth by early Ras1 activation to maintain polarity across generations

**DOI:** 10.1101/2025.10.10.681675

**Authors:** Walker H. Vickers, Maitreyi E. Das

**Author notes:** Funding source: The following work was supported through an NIH/NIGMS R01GM136847 award.

## Abstract

As polarized cells divide, daughter cells must re-establish polarity at the correct site to maintain their shape. Fission yeast cells are bipolar, however growth initiates first at the old end of the cell that was inherited from the mother. This dominance of the old end is dependent on that end having grown in the previous generation, suggesting cells must be able to recognize when an end has a previous history of growth. We find that the proteins Rax1/2 act as markers for memory of growth by localizing to growing cell ends and remaining through division. Rax1/2 then promote activation of Ras1-GTPase at dividing cell ends by recruitment of its GEF Efc25. Localized Ras1 activity at the old ends ensures timely activation of the Rho GTPase Cdc42 to initiate polarized growth, providing an advantage to the old end. Through this work, we demonstrate the role of a Ras-GTPase in positioning the site of polarization and maintaining cell shape across multiple generations.

## Introduction

The importance of structure in determining function is a fundamental principle throughout biology, including proteins, cells, and multicellular organisms. The structure of a cell is established by polarized growth regulated by spatial cues that remodel the cytoskeleton to alter cell shape (Drubin and Nelson, 1996). Polarity plays a role in many processes including cell migration, epithelial morphogenesis, and neural development (Etienne-Manneville, 2004; Zegers and Friedl, 2014; Pichaud et al., 2019; Gu et al., 2023). Disruption of polarity also contributes to cancer development and metastasis (Fomicheva et al., 2020; Peglion and Etienne-Manneville, 2023). Because of its importance, the establishment of polarity requires careful regulation. However, cells not only need to initially establish proper polarity, but also maintain it over time and across multiple generations. Many cells change or lose polarization during cell division, requiring newly formed daughter cells to repolarize to initiate growth at the correct site and form their proper shape. Therefore, investigating the persistence and inheritance of polarity is important to better understand its regulation and proper function.

Many of the major regulators of polarity are highly conserved in eukaryotes, underscoring their importance (Chiou et al., 2017). This includes GTPases such as Cdc42, Rac, and Rho, which act as molecular switches in the regulation of many processes throughout the cell (Johnson, 1999; Nobes and Hall, 1999; Etienne-Manneville, 2004; Mosaddeghzadeh and Ahmadian, 2021). GTPases are activated by binding of GTP, which is promoted by guanine nucleotide exchange factors (GEFs), and deactivated when the hydrolysis of GTP to GDP is facilitated by GTPase-activating proteins (GAPs) (Bos et al., 2007). Mammalian systems have a wide diversity of GEFs and GAPs, with many regulating the same GTPase, making study of the details of their role in polarization difficult (Etienne-Manneville, 2004; Cook et al., 2014; Mosaddeghzadeh and Ahmadian, 2021). By taking advantage of the conservation of the major regulators in simpler eukaryotes such as yeast, we can build a better understanding of the underlying principles in polarity regulation and use it to guide research into higher eukaryotes. In particular, the fission yeast *Schizosaccharomyces pombe* is an excellent model system to study polarity, as it’s rod-like shape and the ability to transition from monopolar to bipolar growth allow for detailed quantification of changes in polarized growth.

While fission yeast cells are bipolar, they first grow only from the old end which is inherited from the mother cell. Growth from the new end formed by the division site is delayed until cells are approximately 9.5 µm in length, when bipolar growth begins by a process known as new-end take-off (NETO) (Mitchison and Nurse, 1985). Polarized growth is initiated by activation of Cdc42 by its GEFs, Gef1 and Scd1 (Chang et al., 1994; Coll et al., 2003; Kelly and Nurse, 2011; Hercyk et al., 2019). The two ends compete for Cdc42 activity as evidenced by its anti-correlated oscillatory dynamics (Das et al., 2012; Xu and Jilkine, 2018; Khalili et al., 2020). The old end initially outcompetes the new end and continues to have an advantage, maintaining a higher growth rate and increased Cdc42 activity even during bipolar growth (Das et al., 2012). This bias toward initiating growth at the old end is despite Cdc42 being primarily active at the division site (the future new end) and deactivated at the old ends during division (Tatebe et al., 2008; Wei et al., 2016). This suggests a mechanism of old-end dominance that promotes activation of Cdc42 and initiation of growth at that end. One factor important for old-end dominance is a previous history of growth. A cell end with a history of growth in one generation will grow in the next. This is true even for most cell polarity mutants (Huisman and Brunner, 2011). Conversely, the daughters of monopolar *gef1Δ* cells which inherit a non-growing end are precociously bipolar, losing old-end dominance (Hercyk et al., 2019). In another category of monopolar mutants such as *tea1Δ*, the old end in daughter cells with no history of growth fails and instead growth occurs only at the new end (Niccoli et al., 2003). The cell must therefore have a memory of growth that allows it to recognize when an end has grown in a previous generation.

Polarity markers that act as a memory across multiple generations have been identified in the regulation of bud-site selection in *Saccharomyces cerevisiae*. The proteins Rax1 and Rax2 localize to the mother-bud neck and then remain at the former division site, marking bud scars through multiple cell cycles and preventing re-budding at the same site (Chen et al., 2000; Fujita et al., 2004; Meitinger et al., 2014). In diploid cells they promote a bipolar budding pattern through interaction with polarity landmarks Bud8/9, leading to activation of the Ras-like GTPase Rsr1 (Kang et al., 2004; Krappmann et al., 2007; Bi and Park, 2012). Rsr1 then promotes activation of Cdc42 through its GEF Cdc24 (Kang et al., 2010; Lee et al., 2015). Homologs of Rax1/2 in fission yeast have been identified and shown to play a role in regulation of polarity (Choi et al., 2006; Johnson, 2019), however their specific function and the mechanism of their regulation in fission yeast are not well understood.

We find that the proteins Rax1/2 act as a memory of growth in fission yeast, localizing to cell ends in a growth-dependent manner and acting as a polarity landmark in newly-divided daughter cells. When the Rax1/2 complex is disrupted, old-end dominance decreases and competition for Cdc42 activation between the two ends shifts to become more equal. The Rax1/2 complex promotes old-end dominance by recruitment of the Ras1-GTPase GEF Efc25 to dividing cell ends, increasing localized Ras1 activation. Ras1 has been shown to act as a functional homolog to Rsr1, promoting activation of Cdc42 through its GEF Scd1 (Lamas et al., 2020). Here we show the role of Rax1/2 and Ras1 in positioning the site of polarized growth initiation to maintain cell shape from one generation to the next.

## Results

### Rax1 regulates the timing and rate of new-end growth

Previous reports on Rax1/2 in fission yeast found an increase in bipolar cells and premature NETO in both *rax2Δ* and *rax1Δ* cells (Choi et al., 2006; Johnson, 2019). These findings are consistent with what we would expect of a memory of growth factor, as its loss should decrease old-end dominance. To determine the role of Rax1/2 in memory of growth, we deleted *rax1* (SPAC23G3.05c) and investigated cell polarity. Consistent with the previously described work, *rax1Δ* increases the percentage of cells with bipolar growth as shown by calcofluor staining (Fig 1 A,B). We then measured the timing of NETO and the growth rate of each end using timelapse imaging. Cells lacking *rax1* undergo premature NETO, with growth initiating at the new end earlier in *rax1Δ* cells than *rax1^+^* (Fig 1 C,D). While the total growth rate of the cell does not change, growth at the old end decreases in *rax1Δ* while the new-end growth rate increases, demonstrating a loss of old-end dominance (Fig 1 E-G). However, the dominance of the old end is not fully abolished with the deletion of *rax1*, as the old end continues to initiate growth first and grow at a higher rate than the new end (Fig 1 E). Instead, loss of *rax1* decreases the advantage of the old end, allowing the new-end to better compete for growth.

**Figure 1.**
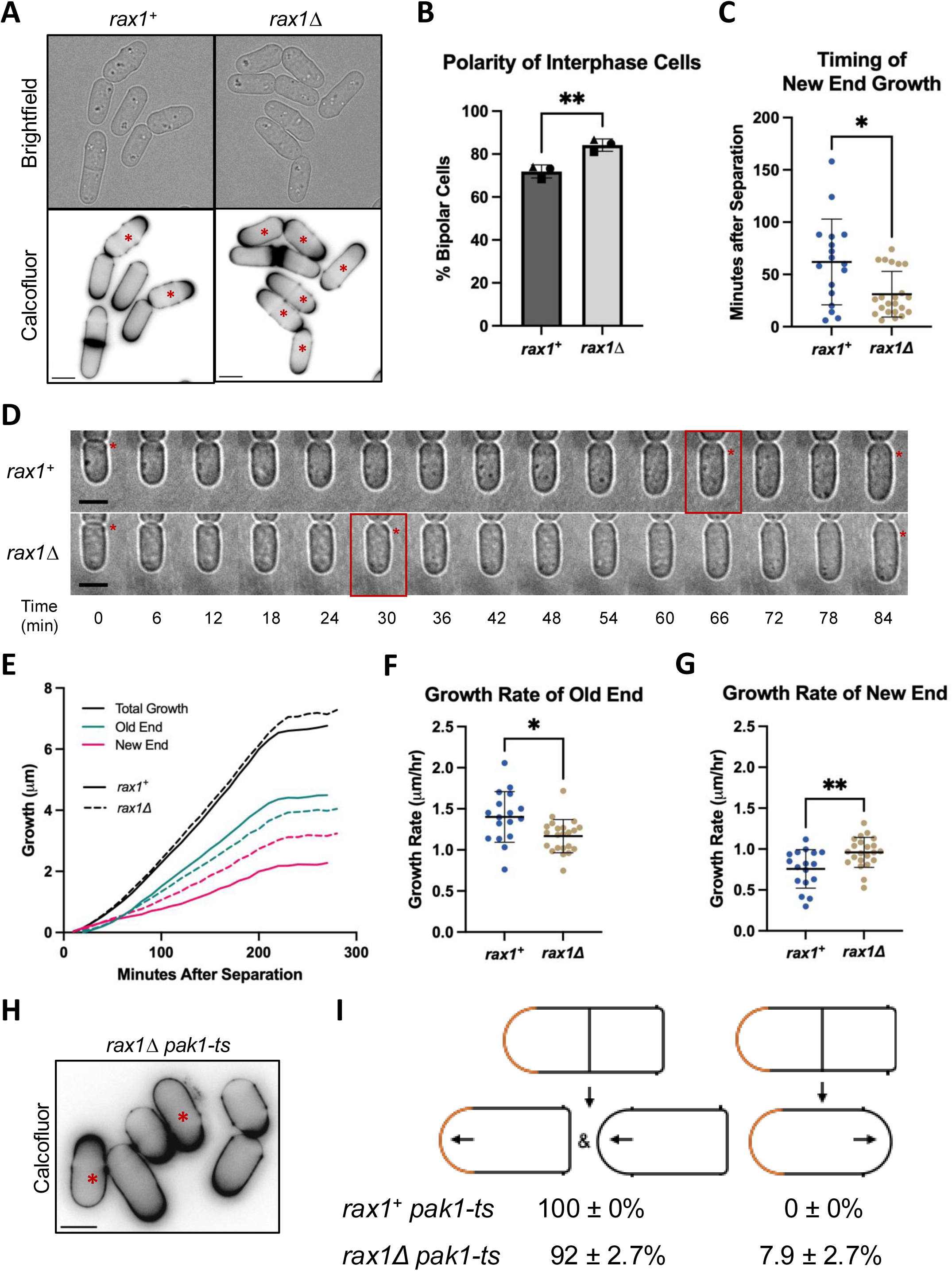
Rax1 promotes old-end dominance and delays new-end growth. **(A)** Polarized growth of *rax1^+^* and *rax1Δ* cells stained with calcofluor white. Asterisks mark bipolar cells. **(B)** Percentage of interphase cells displaying bipolar growth as determined by calcofluor staining. Each point corresponds to an independent experiment (n ≥ 183 cells, N = 3). **(C)** Quantification of the timing of onset of new-end growth in *rax1^+^* and *rax1Δ* cells. Each point represents a cell from one of three independent experiments (n ≥ 17) **(D)** Montage of brightfield images of *rax1^+^*and *rax1Δ* cells. Time ‘0’ marks the onset of cell separation, red box marks onset of new-end growth, asterisks mark the birth scar of the new end. **(E)** Mean increase in old end, new end, and total cell length over the cell cycle in *rax1^+^* and *rax1Δ* cells. **(F,G)** Average growth rate after onset of new-end growth at the old end (F) and new end (G) of *rax1^+^* and *rax1Δ* cells. Each point represents a cell imaged in one of three independent experiments. **(H)** Calcofluor staining of *rax1Δ pak1-ts* cells. Asterisks mark cells with monopolar growth at the new end that inherited an old end with a history of growth. **(I)** Growth patterns of cells with and without *rax1* in *pak1-ts* mutants. Orange marks previously growing ends, arrows mark new growth. Percentages were determined by calcofluor staining and are mean ± s.d. from three independent experiments (n ≥ 158 cells, N = 3). Scale bars are 5 µm. Error bars are s.d. Significance was determined using an unpaired, two-tailed t-test with Welch’s correction. * = p < 0.05, ** = p < 0.01.

This decreased old-end dominance was also apparent when *rax1Δ* was combined with deletions of other polarity regulators, including mutants with increased monopolarity such as *gef1Δ* and *scd2Δ*. In most cases, deletion of *rax1* increases the percentage of bipolar cells relative to the mutant background (Table S1). While loss of *rax1* fails to rescue bipolar growth in the strictly monopolar *orb2-34* (*pak1-ts*) mutant, we did observe a change in growth pattern. Cells with the *pak1-ts* allele fail to become bipolar. The daughter cell that inherits the previously growing end grows only from the old end, while the daughter that inherits a non-growing end instead initiates growth at the new end (Kim et al., 2003). However, when observing calcofluor stained *rax1Δ pak1-ts* cells, we identified a small percentage of cells that appear to have inherited an end with a previous history of growth yet fail to re-establish growth at that site. Instead the daughter cells grow from the former division site as if they inherited a non-growing end. (Fig 1 H,I). This supports the hypothesis that Rax1 acts as the memory of growth and deletion of *rax1* decreases the ability of the cell to recognize previously growing ends.

### Rax1/2 localize to cell ends in a growth-dependent manner

The function of a memory-of-growth marker requires that its localization to the ends be dependent on growth. It must then remain stable at those sites of growth through division. Both Rax1 and Rax2 have been reported to localize to cell ends, including in dividing cells (Choi et al., 2006; Johnson, 2019). We visualized Rax1 and Rax2 by tagging with an N-terminal mNeonGreen (mNG) or a C-terminal GFP respectively. Both appear as puncta along the cell membrane clustered at cell ends in both growing and dividing cells (Fig 2 A). mNG-Rax1 and Rax2-GFP also localize in a non-constricting ring at the division site (Fig S1 A). mNG-Rax1 appears to be less abundant than Rax2-GFP, as it was more difficult to image and required higher laser power, though a direct comparison was not made due to the difference in fluorophore. Consistent with their *S. cerevisiae* homologs and previous reports (Kang et al., 2004; Johnson, 2019), localization of Rax2 to the cell ends is dependent on Rax1, with Rax2-GFP failing to localize to the cell ends in a *rax1Δ* background (Fig S1 B).

**Figure 2.**
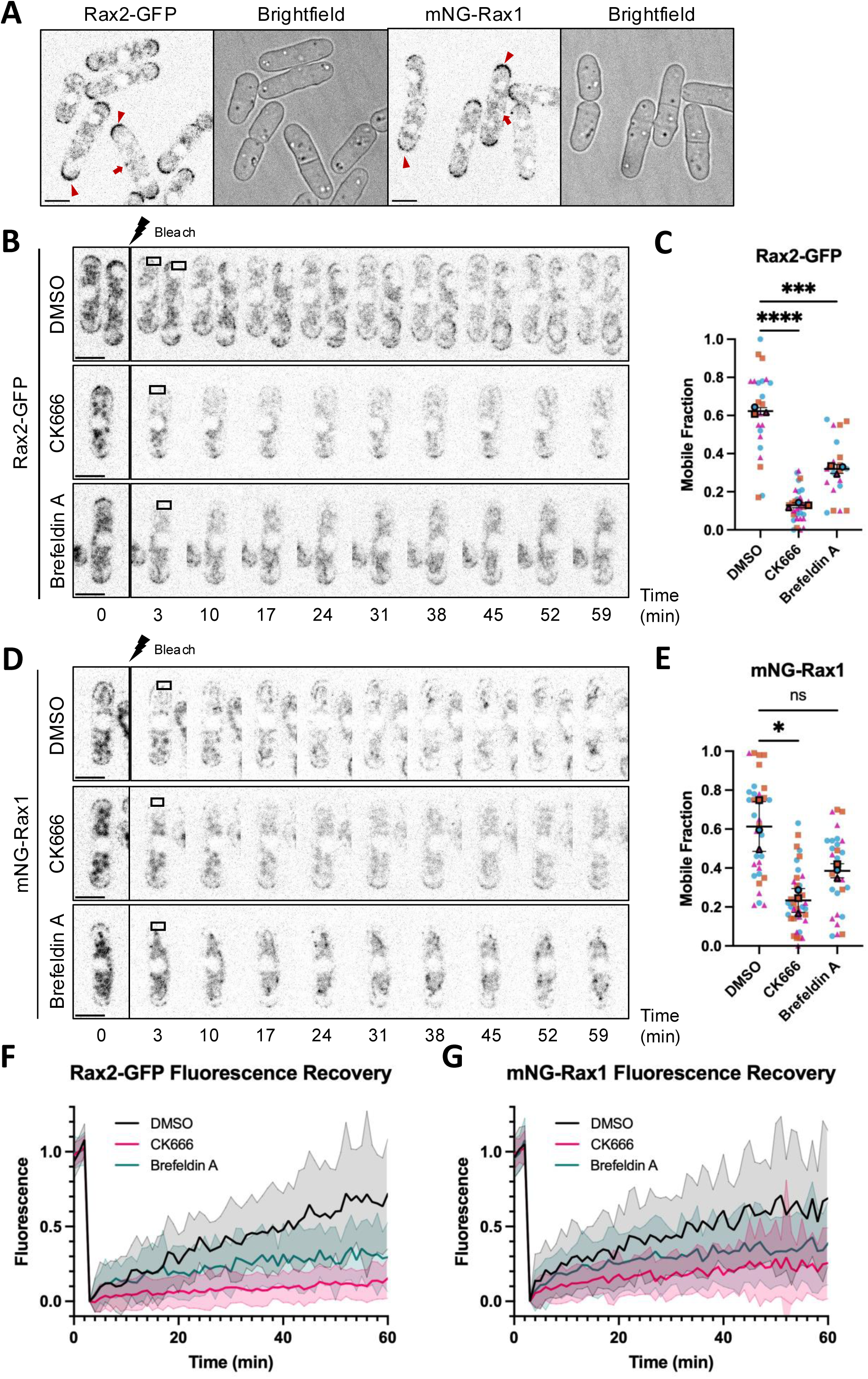
Localization of Rax1/2 to cell ends is growth-dependent. **(A)** Localization of Rax2-GFP and mNG-Rax1 to cell ends. Arrowheads show localization to the old ends of dividing cells and newly formed daughter cells, while arrows mark division site localization. **(B)** Montage showing fluorescence recovery after photobleaching of Rax2-GFP in cells treated with DMSO (1%), CK666 (100 µM), or Brefeldin A (100 µg/mL). **(C)** Quantification of the mobile fraction of Rax2-GFP fluorescence recovery after 60 minutes. Outlined points represent the mean from one independent experiment with small points corresponding to individual cells (n ≥ 6 cells, N = 3). **(D)** Montage showing fluorescence recovery after photobleaching of mNG-Rax1 in cells treated with DMSO (1%), CK666 (100 µM), or Brefeldin A (100 µg/mL). **(E)** Quantification of the mobile fraction of mNG-Rax1 fluorescence recovery after 60 minutes. Outlined points represent the mean from one independent experiment, with small points corresponding to individual cells (n ≥ 7 cells, N = 3). **(F,G)** Mean and s.d. of normalized fluorescence recovery of Rax2-GFP (F) and mNG-Rax1 (G) over time (n ≥ 20 cells). Scale bars are 5 µm. Error bars are s.d. Significance determined by one-way ANOVA followed by Dunnett’s multiple comparisons test, * = p < 0.05, *** = p < 0.001, **** = p < 0.0001, ns = not significant.

To confirm that cell end localization of the Rax1/2 complex is growth-dependent we performed fluorescence recovery after photobleaching (FRAP) at cell ends. Growth was inhibited by treatment with CK666, an Arp2/3 inhibitor (Nolen et al., 2009), or Brefeldin A, an ER-to-Golgi transport inhibitor (Turi et al., 1994), with DMSO as a control. Both mNG-Rax1 and Rax2-GFP display fluorescence recovery at the ends of DMSO treated interphase cells consistent with a slow accumulation during growth (Fig 2 B,D,F,G). When growth was arrested with CK666 or Brefeldin A fluorescence recovery was lost, with a significant decrease in the mobile fraction of Rax2-GFP in both conditions (Fig 2 C, F). mNG-Rax1 shows a similar trend though with a smaller effect, likely due to increased bleaching during image acquisition and its lower abundance (Fig 2 E,G). We also examined fluorescence recovery of Rax1/2 at dividing cell ends in the DMSO control, as growth arrests during division (Ray et al., 2010). Rax2-GFP and mNG-Rax1 both show decreased fluorescence recovery in dividing cells, further supporting the growth-dependent localization of Rax1/2 (Fig S1 C,D).

### Rax1 promotes activation of Cdc42 at the old ends of separating cells

While our data show that Rax1 and Rax2 act as memory of growth markers, we wanted to identify the mechanism for how they promote old-end dominance. Therefore, we examined the localization of the major regulator of polarized growth, Cdc42. The dominance of the old end is evident in the competition for active Cdc42, with the old end maintaining higher levels of Cdc42 activation until late in the cell cycle, after bipolar growth has initiated (Das et al., 2012). When visualizing Cdc42-GTP levels using the fluorescent probe CRIB-3xGFP (Tatebe et al., 2008), a higher fraction of the total end intensity of CRIB-3xGFP is present at the old end than the new end. However, *rax1Δ* increases the proportion of Cdc42 activity present at the new end, leading to a more equal distribution between the two ends (Fig 3 A,B). We also observe this shift in distribution towards the new end in the localization of the main GEF for Cdc42, Scd1, and its scaffold, Scd2 (Fig 3 A,C,D). Interestingly, while the growth rate at the old end of *rax1Δ* cells decreases along with the increase in new-end growth, there is no significant change in the level of Cdc42-GTP or its activators at the old end. Instead, more equal distribution between the two ends is primarily due to an increase at the new end (Fig S2). Growth rate is therefore not directly determined by the amount of Cdc42-GTP but must be regulated by another limiting factor.

**Figure 3.**
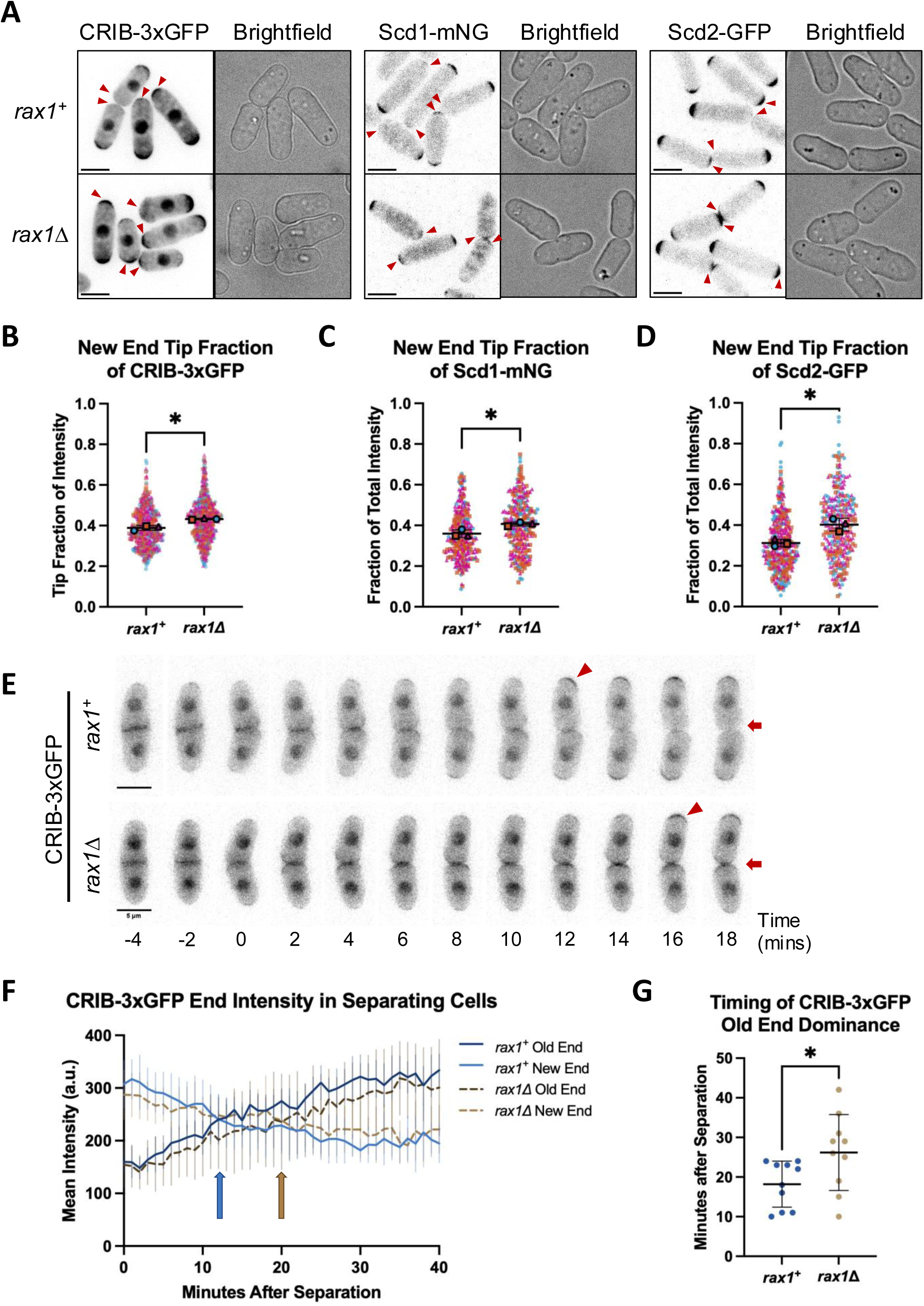
Loss of *rax1* decreases old-end dominance by delaying Cdc42 activation at that end during cell separation. **(A)** Localization of CRIB-3xGFP, Scd1-mNG, and Scd2-GFP at the ends of *rax1^+^* and *rax1Δ* cells. Images are sum projections of z series. Arrowheads mark new ends. **(B-D)** Quantification of the fraction of total tip intensity of CRIB-3xGFP (B), Scd1-mNG (C), and Scd2-GFP (D) at the new ends of *rax1^+^* and *rax1Δ* cells. Outlined points represent the mean from one independent experiment, with small points corresponding to individual cells (n ≥ 54 cells, N = 3). **(E)** Montage of CRIB-3xGFP in separating *rax1^+^* and *rax1Δ* cells. Arrowheads mark localization of CRIB-3xGFP at old ends, while arrows mark the new ends. Time ‘0’ is onset of cell separation. **(F)** Quantification of mean and s.d. of CRIB-3xGFP intensity over time at the old and new ends of separating *rax1^+^* and *rax1Δ* cells (n = 10). Arrows mark the point where old- end intensity becomes higher than new-end intensity. **(G)** Quantification of the timepoint at which old-end CRIB-3xGFP intensity becomes higher than new-end intensity. Data points correspond to individual cells. Scale bars are 5 µm. Error bars are s.d. Significance was determined using an unpaired, two-tailed t-test with Welch’s correction. * = p < 0.05.

To further understand how *rax1Δ* cells decrease in old-end dominance and initiate early new-end growth, we also measured the dynamics of Cdc42 activity in newly separating daughter cells by timelapse imaging of CRIB-3xGFP. In the wildtype, Cdc42 is active at the division site until just before cell separation, when Cdc42 activity shifts to the old end and leaves the recently formed new end. However, CRIB-3xGFP remains longer at the new end of *rax1Δ* cells, with a delay in accumulation at the old end (Fig 3 E). The timepoint where the old end begins to outcompete the new end, when CRIB-3xGFP intensity at the old end becomes higher than at the new end, is delayed by ∼8 minutes in *rax1Δ* (Fig 3 F, gold arrow) compared to *rax1^+^* (Fig 3 F, blue arrow). We find that in *rax1^+^*cells CRIB-3xGFP intensity at the old end overcomes new end intensity 18 minutes after cell separation while in *rax11* cells this occurs after 26 minutes (Fig 3 G). This delay in old end localization is also present in Scd1-mNG and Scd2-GFP (Fig S3). These results show that Rax1 promotes timely re-initiation of Cdc42 activation at the cell ends following division.

### Ras1-GTPase activity at the ends of dividing cells is dependent on Rax1

Next, we investigated how the Rax1/2 complex regulates Cdc42 activity. In *S. cerevisiae*, polarized activation of Cdc42 downstream of Rax1/2 is through the activity of a Ras-like GTPase, Rsr1 (Kang et al., 2010; Lee et al., 2015). The Rax1/2 complex promotes localization of polarity landmark proteins Bud8/9, which recruit the GEF of Rsr1, Bud5 (Kang et al., 2004; Krappmann et al., 2007). Active Rsr1 then interacts with the GEF Cdc24, which activates Cdc42 (Chiou et al., 2017). While homologs of Bud8/9 have not been identified in fission yeast, the Ras-GTPase in fission yeast, Ras1, has been found to promote polarized activation of Cdc42 through interaction with Scd1, acting in a functionally similar pathway to Rsr1 (Lamas et al., 2020; Chang et al., 1994). We therefore hypothesized that Rax1/2 may regulate Cdc42 through Ras1. We visualized Ras1 activity using the fluorescent probe RasAct^GFP^, which binds to the active form Ras1-GTP (Merlini et al., 2018). In contrast to Cdc42, Ras1 is highly active at the ends of dividing cells as well as the division site (Fig 4 A,D). Without *rax1* RasAct^GFP^ no longer localizes to dividing cell ends, instead showing increased intensity at the division site (Fig 4 A-E). In timelapse imaging of dividing cells, Ras1 activity increases at the cell ends until it peaks right before cell separation (Fig 4 D,E, red arrowhead). However, in the *rax1Δ* activation of Ras1 instead peaks at the division site, with Ras1 primarily active at the new end right after cell separation (Fig 4 D,E, blue arrowhead). Interestingly, this change in Ras1 activity is limited to division, as RasAct^GFP^ localizes to the ends of interphase cells at similar intensities in both *rax1Δ* and *rax1^+^* cells (Fig 4 F,G). If Ras1 activation at dividing cell ends is dependent on Rax1, then there should be no active Ras1 at ends without a previous history of growth. We therefore examined RasAct^GFP^ localization in monopolar cells, specifically in a *tea1Δ* strain. Tea1 is delivered to the cell ends through microtubules and is required for the initiation of a second site of growth (Kim et al., 2003; Niccoli et al., 2003). Consistent with our hypothesis, Ras1 activity is limited to the previously growing end of dividing *tea1Δ* cells and is absent from the non-growing end (Fig 4 H).

**Figure 4.**
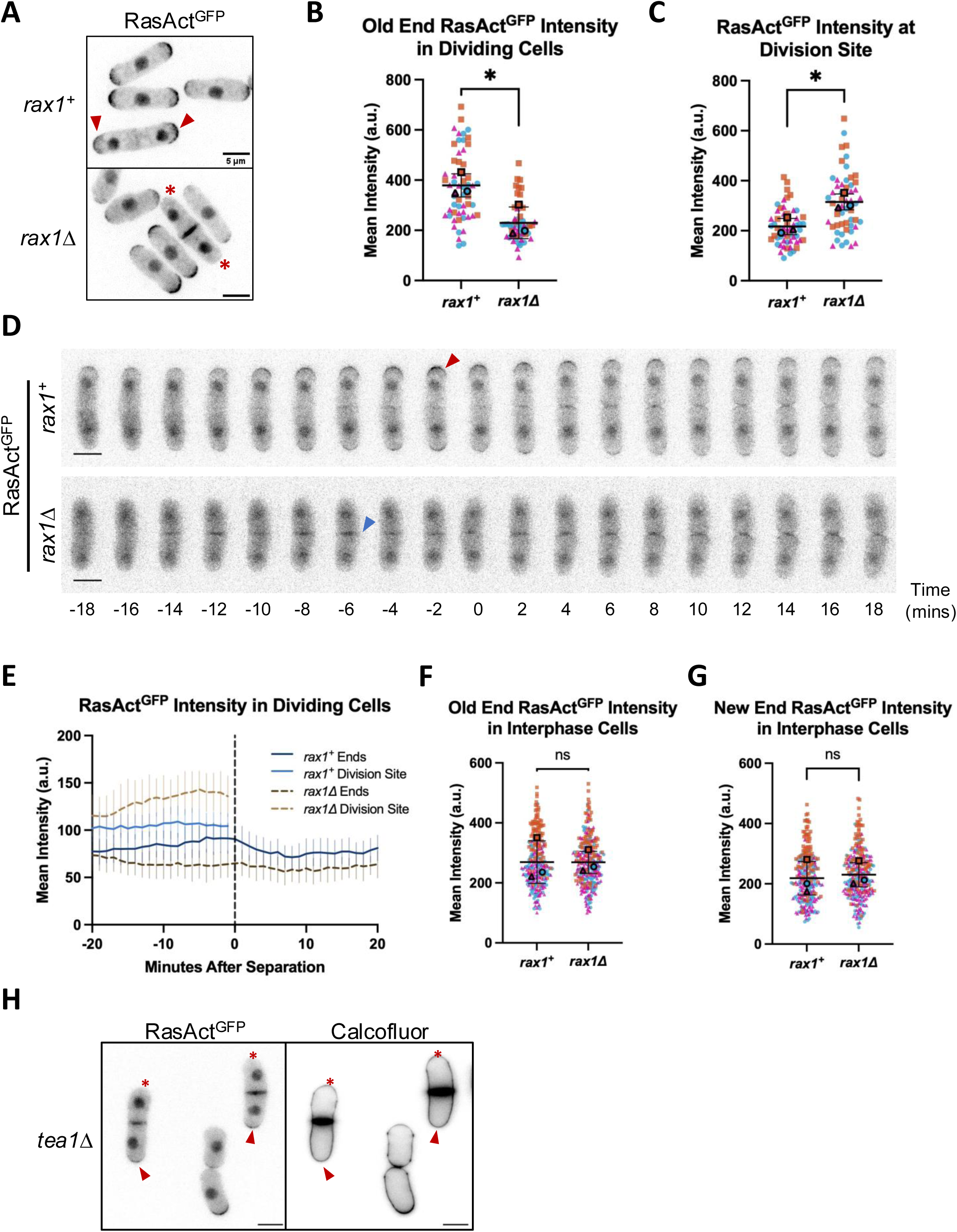
Rax1 promotes activation of Ras1 at the ends of dividing cells. **(A)** Localization of RasAct^GFP^ in *rax1^+^*and *rax1Δ* cells. Images are sum projections of z series. Arrowheads mark Ras1 activity at the ends of dividing cells, while asterisks mark dividing cell ends without Ras1 activity. **(B,C)** Quantification of RasAct^GFP^ intensity at the ends (B) and division site (C) of dividing cells. Outlined points represent the mean from one independent experiment, with small points corresponding to individual cells. (n ≥ 15 cells, N = 3) **(D)** Montage of RasAct^GFP^ in separating *rax1^+^*and *rax1Δ* cells. The red arrowhead marks the peak intensity of RasAct^GFP^ at the ends of *rax1^+^* cells, and the blue arrowhead marks the peak intensity at the division site in *rax1Δ* cells. Time ‘0’ is onset of cell separation. **(E)** Quantification of the mean and s.d. of RasAct^GFP^ intensity over time in *rax1^+^*and *rax1Δ* cells mid-separation (n = 10). **(F,G)** Quantification of mean RasAct^GFP^ intensity at the old ends (F) and new ends (G) of interphase *rax1^+^* and *rax1Δ* cells. Outlined points represent the mean from one independent experiment, with small points corresponding to individual cells. (n ≥ 78, N = 3) **(H)** Localization of RasAct^GFP^ in *tea1Δ* cells stained with calcofluor white. Arrowheads mark dividing cells where RasAct^GFP^ is localized to the end that shows evidence of growth in the previous generation, with asterisks marking non-growing ends lacking RasAct^GFP^. The RasAct^GFP^ image is a sum projection of a z series. Scale bars are 5 µm. Error bars are s.d. Significance was determined using an unpaired, two-tailed t-test with Welch’s correction. * = p < 0.05, ns = not significant.

### Active Ras1 promotes old-end dominance

While our results show that Rax1 regulates the localization of Ras1 activity during division, our next step was to determine whether this loss of Ras1-GTP from dividing cell ends is responsible for the loss of old-end dominance observed in *rax1Δ* cells. To address this, we investigated the polarity phenotype of cells lacking active Ras1. While *ras1Δ* cells are sterile, deletion of the Ras1 GEF Efc25 removes Ras1 activation during mitotic growth but does not have mating defects (Fukui et al., 1986; Papadaki et al., 2002; Merlini et al., 2018). Thus, we crossed *efc25Δ* with *rax1Δ* and looked at the bipolarity of the single and double deletions with calcofluor staining. Despite being much shorter and wider with clear shape defects, *efc25Δ* cells maintain a high percentage of bipolarity (Fig 5 A,B). However, unlike previously tested deletions, loss of *rax1* in an *efc25Δ* background does not increase bipolarity compared to *rax1^+^* (Fig 5 B). This epistatic interaction between *efc25Δ* and *rax1Δ* indicates that activation of Ras1, likely through Efc25, is downstream of Rax1/2.

**Figure 5.**
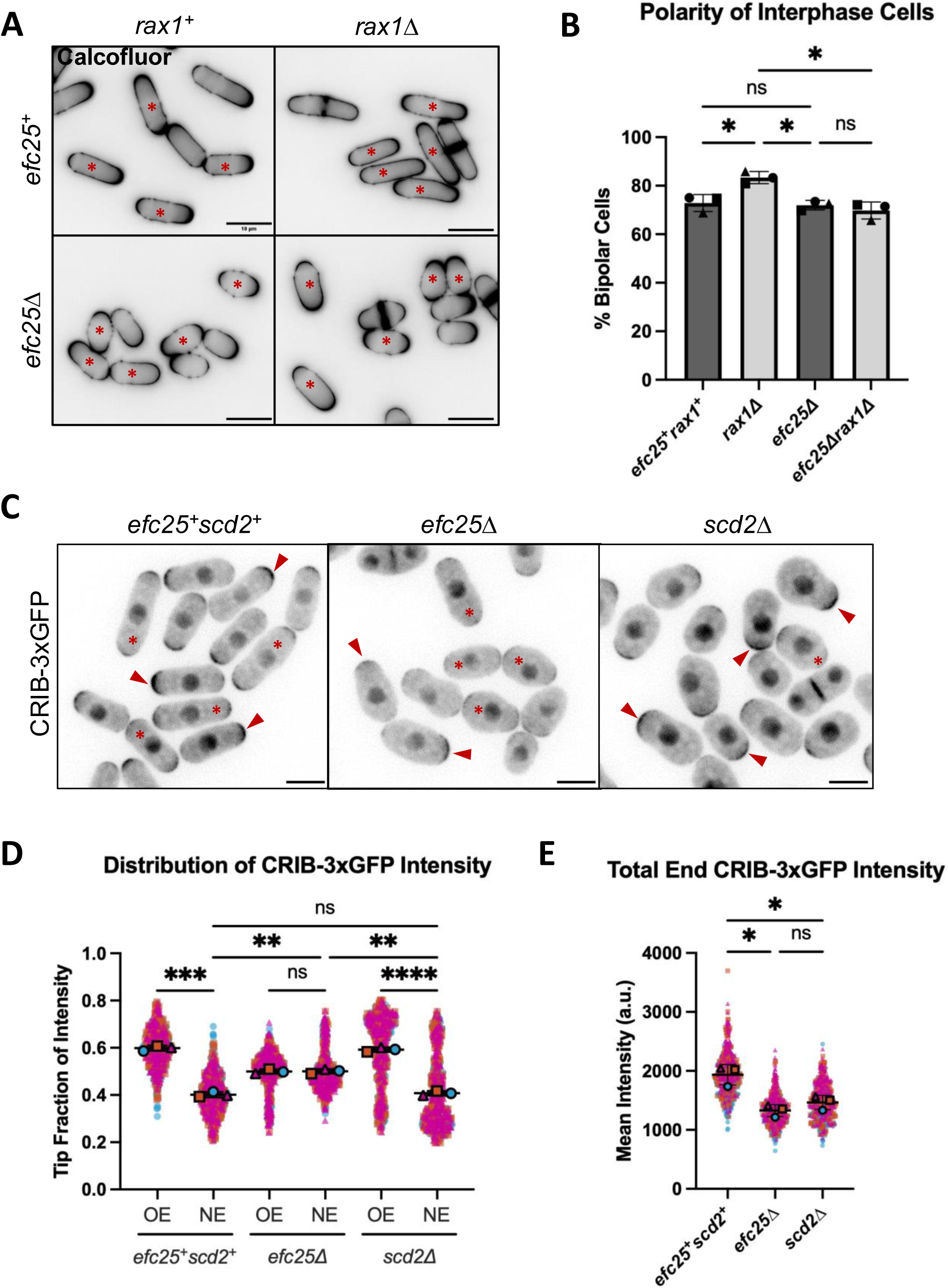
Activation of Ras1 by its GEF Efc25 promotes Rax1-mediated old-end dominance. **(A)** Polarized growth of cells stained with calcofluor white in the indicated strains. Asterisks mark bipolar cells. Scale bars are 10 µm. **(B)** Percentage of interphase cells displaying bipolar growth as determined by calcofluor staining. Each point corresponds to the mean of an independent experiment (n ≥ 174, N = 3). **(C)** Localization of CRIB-3xGFP. Images are sum projections of z series. Asterisks mark cells with similar levels of CRIB-3xGFP at both ends, while arrowheads mark examples of cells with asymmetric CRIB-3xGFP distribution. Scale bars are 5 µm. **(D)** Quantification of the fraction of total CRIB-3xGFP tip intensity at the old end (OE) and new end (NE). Outlined points represent the mean from one independent experiment, with small points corresponding to individual cells (n ≥ 147 cells, N = 3). **(E)** Quantification of the total CRIB-3xGFP intensity at the cell ends. Outlined points represent the mean from one independent experiment, with small points corresponding to individual cells. Error bars are s.d. Significance determined by one-way ANOVA followed by Dunnett’s multiple comparisons test, * = p < 0.05, ** = p < 0.01, *** = p < 0.001, **** = p < 0.0001, ns = not significant.

If active Ras1 promotes old-end dominance, then cells lacking Ras1 activity should show similar phenotypes to those observed in *rax1Δ*. Therefore, we investigated the distribution of Cdc42 activity in *efc25Δ* cells and found that CRIB-3xGFP intensity is more evenly divided between the two ends than the wildtype (Fig 5 C,D). Because Ras1 promotes Scd1 activity at the cell ends, cells lacking active Ras1 also have less Cdc42 activity, similar to the deletion of *scd2*, the scaffold for Scd1 (Lamas et al., 2020) (Fig 5 E). Despite similar decreases in the total active Cdc42 at cell ends, *scd2Δ* cells display a highly asymmetric distribution of CRIB-3xGFP, indicating that the increased fraction of Cdc42 activity at the new end is specific to *efc25Δ* and not an artifact of less available Cdc42-GTP (Fig 5 C-E). Therefore, Ras1 activity does promote old-end dominance, as this dominance is decreased when Ras1 activation is lost.

### Efc25 localization to dividing cell ends is dependent on Rax1

Because Efc25 is the main GEF of Ras1 during mitotic growth and is required for Ras1’s role in morphogenesis (Tratner et al., 1997; Papadaki et al., 2002), we hypothesized that the Rax1/2 complex regulates activation of Ras1 through Efc25. Therefore, we examined localization of Efc25 with and without *rax1*. We first imaged Efc25 tagged with GFP at the C-terminus expressed under its native promoter. While in dividing cells Efc25-GFP appears primarily at the division site, we also observe faint localization at the cell ends (Fig 6 A). When measuring Efc25-GFP intensity along the cell cortex, we find that in *rax1^+^*cells Efc25-GFP intensity peaks at the tips of the cell ends, and that this peak is severely diminished in *rax11* cells (Fig 6 B). We also observed a trend in increasing Efc25-GFP intensity at the division site of *rax11* cells, however the change was not statistically significant (Fig 6 C, p=0.0908). The low abundance of Efc25 and biological noise may have obscured the signal difference that we observed between these strains. To more clearly visualize Efc25 localization, we tagged it with an N-terminal mNeonGreen and expressed it under the *nmt41* thiamine-inducible promoter (Basi et al., 1993). We then grew cells in minimal media lacking thiamine and imaged mNG-Efc25. When overexpressed, mNG-Efc25 clearly localizes to dividing cell ends as well as the division site in *rax1^+^* cells (Fig 6 D). Without *rax1*, mNG-Efc25 no longer accumulates at dividing cell ends and instead increases at the division site, consistent with the trends seen with the endogenously tagged Efc25-GFP (Fig 6 D-F). Interestingly, total intensity of both Efc25-GFP and mNG-Efc25 is higher in *rax1Δ* cells than in *rax1^+^*, suggesting that Rax1 not only regulates Efc25 localization but also its protein levels (Fig 6 G,H). However, it appears that it is the change of distribution of Efc25 between the ends and the division site that is important for old-end dominance and not the amount of protein, as overexpression of mNG-Efc25 does not affect the percentage of bipolar cells in *rax1^+^* or *rax1Δ* cells (Fig 6 I).

**Figure 6.**
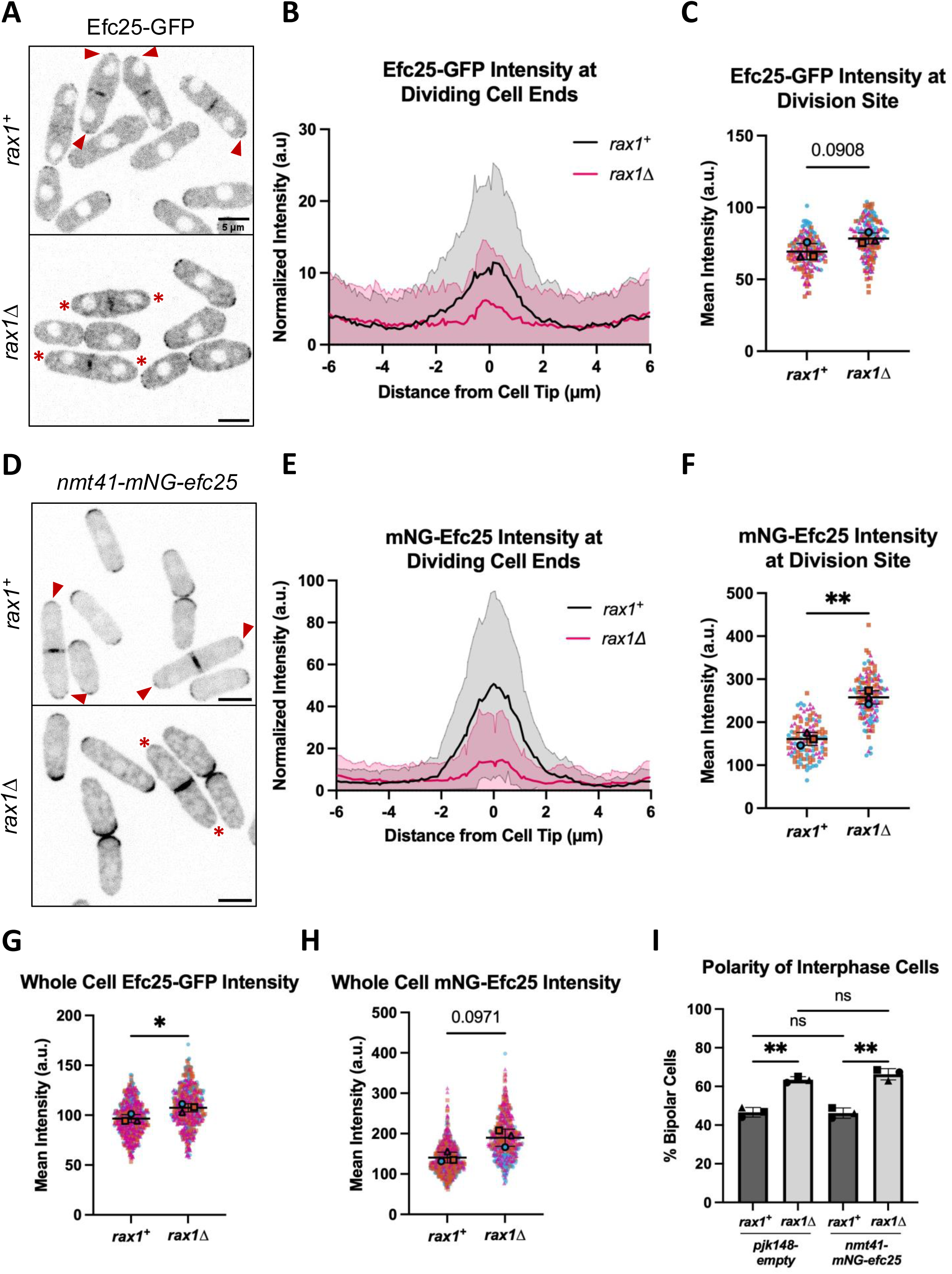
Rax1 promotes localization of Efc25 to dividing cell ends. **(A)** Localization of Efc25-GFP in *rax1^+^* and *rax1Δ* cells. Images are sum projections of the middle 4 planes of a z series. Arrowheads mark Efc25-GFP present at the ends of dividing cells, while asterisks mark dividing cell ends without Efc25-GFP. **(B)** Plot profile of the mean and s.d. of normalized Efc25-GFP intensity along the cell cortex in dividing cells (n ≥ 157). Intensity was normalized to that at the cell sides. **(C)** Quantification of Efc25-GFP intensity at the division site. Outlined points represent the mean from one independent experiment, with small points corresponding to individual cells. Significance was determined using an unpaired, two-tailed t-test with Welch’s correction. (n ≥ 43, N = 3). **(D)** Localization of mNG-Efc25 expressed from the *nmt41* promoter in *rax1^+^* and *rax1Δ* cells grown in EMM lacking thiamine. Images are sum projections of the middle 4 planes of a z series. Arrowheads mark mNG-Efc25 present at the ends of dividing cells, while asterisks mark dividing cell ends without mNG-Efc25. **(E)** Plot profile of the mean and s.d. of normalized mNG-Efc25 intensity along the cell cortex in dividing cells (n ≥ 115). Intensity was normalized to that at the cell sides. **(F)** Quantification of mNG-Efc25 intensity at the division site. Outlined points represent the mean from one independent experiment, with small points corresponding to individual cells. Significance was determined using an unpaired, two-tailed t-test with Welch’s correction. (n ≥ 33, N = 3). **(G,H)** Quantification of whole cell intensity of Efc25-GFP (G) or mNG-Efc25 (H) in *rax1^+^* and *rax1Δ* cells. Outlined points represent the mean from one independent experiment, with small points corresponding to individual cells. Significance was determined using an unpaired, two-tailed t-test with Welch’s correction. (n ≥ 123, N = 3). **(I)** Percentage of interphase cells displaying bipolar growth as determined by calcofluor staining. Each point corresponds to the mean of an independent experiment (n ≥ 308, N = 3). Significance determined by one-way ANOVA followed by Dunnett’s multiple comparisons test. Scale bars are 5 µm. Error bars are s.d. * = p < 0.05, ** = p < 0.01, ns = not significant.

If the Rax1/2 complex promotes old-end dominance through recruitment of Efc25 to dividing cell ends, then we would expect that targeting Efc25 to the ends independently of Rax1/2 would restore old-end dominance in *rax1Δ* cells. We investigated this question by using a GFP-binding protein (GBP) trap (Rothbauer et al., 2008; Chen et al., 2017), tagging Efc25 with GBP and mScarlet and expressing it under the *nmt41* promoter in cells containing Tea1-GFP. We used Tea1-GFP because it localizes to the cell ends and the division site through cell division (Behrens and Nurse, 2002). Tea1-GFP successfully targets GBP-mScarlet-Efc25 to dividing cell ends in *rax1Δ* cells, increasing GBP-mScarlet-Efc25 levels at the cell cortex and reducing cytoplasmic localization (Fig 7 A). When GBP-mScarlet-Efc25 is bound to Tea1-GFP, *rax1Δ* does not increase the percentage of bipolar cells compared to *rax1^+^*, demonstrating that loss of *rax1* has no additional effect when Efc25 is independently targeted to dividing cell ends (Fig 7 B). However, GBP-mScarlet-Efc25 Tea1-GFP cells show increased bipolarity compared to GBP-mScarlet-Efc25 alone, with a similar percentage of bipolarity to *rax1Δ tea1^+^* GBP-mScarlet-Efc25 cells (Fig 7 B). These results are contrary to our prediction that increased Efc25 localization to dividing cell ends would increase old-end dominance. This may be due to Tea1-GFP also increasing GBP-mScarlet-Efc25 localization to the division site and new ends or an effect on GBP-mScarlet-Efc25 activity upon binding to Tea1-GFP.

**Figure 7.**
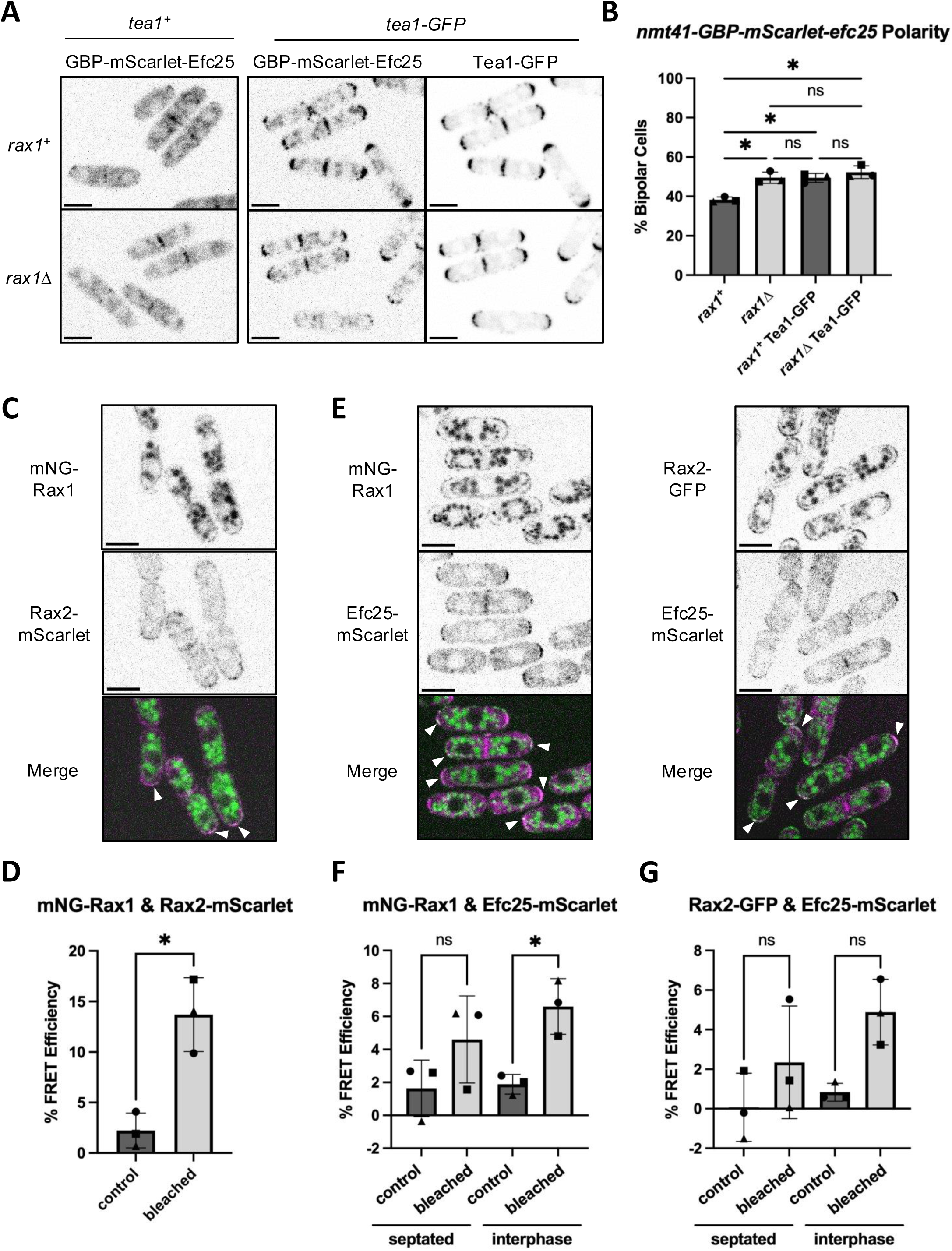
Rax1 and Rax2 form a complex and recruit Efc25 to the cell ends. **(A)** Targeting of GBP-mScarlet-Efc25 to the cell ends and division site by Tea1-GFP. Images are sum projections of the middle 4 planes of a z series. **(B)** Percentage of interphase cells displaying bipolar growth as determined by calcofluor staining. Each point corresponds to the mean of an independent experiment (n ≥ 232, N=3). Significance determined by one-way ANOVA followed by Dunnett’s multiple comparisons test. **(C)** Colocalization of mNG-Rax1 and Rax2-mScarlet. Arrows mark regions of colocalization. **(D)** Quantification of FRET efficiency between mNG-Rax1 and Rax2-mScarlet. Each point represents the mean of an independent experiment (n ≥ 118, N = 3). Significance determined by a paired one-tailed t-test. **(E)** Colocalization of Efc25-mScarlet with mNG-Rax1 or Rax2-GFP. Arrows marks regions of colocalization. **(F,G)** Quantification of FRET efficiency between Efc25-mScarlet and mNG-Rax1 (F) or Rax2-GFP (G). Each point represents the mean of an independent experiment (n ≥ 21, N=3). Significance determined by repeated measures one-way ANOVA followed by Šídák’s multiple comparisons test. % FRET efficiency is the increase in donor fluorescence after photobleaching of the acceptor. Control measurements were taken from unbleached cells in the same field. Scale bars are 5 µm. Error bars are s.d. * = p < 0.05, ns = not significant.

We next investigated whether the recruitment of Efc25 by Rax1/2 is through a direct binding interaction or an indirect mechanism. To determine interaction between Rax1/2 and Efc25 we used Förster resonance energy transfer (FRET) measured by the increase in donor fluorescence after acceptor photobleaching. We first measured FRET efficiency between Rax1 and Rax2 to determine whether they directly bind each other to form a complex. Rax1 and Rax2 have been shown to interact with one another in other yeasts (Kang et al., 2004; Li et al., 2025), so we predicted they would interact in fission yeast as well. Rax1 and Rax2 do colocalize at the cell ends and appear to directly interact with one another, with a FRET efficiency of 13.7%, indicating that they do form a complex (Fig 7 C,D). Next, we measured the FRET efficiency between Efc25-mScarlet with both mNG-Rax1 and Rax2-GFP. Donor fluorescence increased after photobleaching of Efc25-mScarlet with both mNG-Rax1 and Rax2-GFP when measured at the ends of dividing and interphase cells. However, the increase is only statistically significant with mNG-Rax1 at the ends of interphase cells, with a FRET efficiency of 6.61% (Fig 7 E-G). The small amount of Efc25-mScarlet localized to dividing cell ends as well as the relatively low abundance of all three proteins may have impacted the ability to detect interaction through this method. Though further research is required, our results do suggest a potential direct interaction between Rax1 and Efc25.

## Discussion

The delayed bipolar growth pattern in fission yeast occurs because the old end outcompetes the new end for Cdc42 activation. During cytokinesis, the final step in cell division, Cdc42 is inactivated at the cell ends and activated at the division site. However, just as the cells start to separate at the end of division, Cdc42 activity is redirected from the division site to the opposing end of the cell (Tatebe et al., 2008; Wei et al., 2016). Previous reports have shown that Cdc42 activation at the old end is triggered at a certain time after the completion of mitosis and also requires the activation of protein synthesis via the NDR/Orb6 kinase pathway (Rich-Robinson et al., 2021; Verde et al., 1998; Nuñez et al., 2016). The mechanisms that regulate NETO to initiate the second growth site have been studied (Glynn et al., 2001; Martin et al., 2005; Tatebe et al., 2005, 2008; Das et al., 2012; Bohnert and Gould, 2012; Hercyk et al., 2019; Bohnert et al., 2020). However, the mechanisms responsible for old-end dominance that allow it to outcompete the new end for Cdc42 activation are not well understood. The loss of old-end dominance in cells that inherit non-growing ends suggests the presence of a memory of growth (Niccoli et al., 2003; Huisman and Brunner, 2011; Hercyk et al., 2019). Here, we identify the proteins Rax1/2 as memory of growth factors and demonstrate their role in promoting old-end dominance through activation of Ras1.

Rax1/2 homologs have been identified in multiple species of fungi where they appear to have a conserved role in determining the site of polarization. Rax1 and Rax2 interdependently promote bipolar budding in *Saccharomyces cervisiae* and *Yarrowia lipolytica*, and Rax1 has been shown to regulate growth and conidia development in *Aspergillus fumigatus* (Chen et al., 2000; Fujita et al., 2004; Kang et al., 2004; Li et al., 2025; Igbalajobi et al., 2017). Rax2 has also been shown to regulate both bud-site selection and direction of hyphal growth in *Candida albicans* (Gonia et al., 2013). Our results demonstrate that the role of Rax1/2 in positioning the site of polarization is conserved in fission yeast, where they promote initiation of growth first at the old ends of the cell. We also find that similar to how Rax1 and Rax2 mark sites of previous division in *S. cerevisiae* (Chen et al., 2000), in *S. pombe* they localize to the ends in a growth-dependent manner and remain at these ends through cell division, acting as a memory of growth. Consistent with reports in *S. cerevisiae* and *Y. lipolytica* (Kang et al., 2004; Li et al., 2025), we find that Rax1 and Rax2 colocalize and directly interact with each other to form a complex. Their interaction is likely required for their delivery to cell ends, as localization of Rax1 and Rax2 is interdependent. Our data show that without *rax1*, old-end dominance is decreased and cells become precociously bipolar. We find that loss of *rax1* weakens the advantage of the old end across different polarity mutants, including highly monopolar strains, demonstrating the importance of memory of growth to old-end dominance. However, the Rax1/2 complex does not appear to be the only mechanism promoting old-end dominance, as old-end growth still initiates first in *rax1Δ* cells, and deletion of *rax1* only partially rescues bipolar growth in monopolar mutants such as *scd2Δ* and *bud6*Δ (Table S1). This alternative pathway promoting old-end growth is likely Tea1-dependent, since deletion of both *rax2* and *tea1* causes both daughter cells to only grow from their new end (Choi et al., 2006; Johnson, 2019).

Here we show that the Rax1/2 complex promotes old-end dominance by activating Ras1 at dividing cell ends through recruitment of its GEF Efc25. This pathway is similar to the bud-site selection pathway in *S. cerevisiae*, where Rsr1 and its GEF Bud5 are recruited downstream of Rax1/2 (Kang et al., 2004; Bi and Park, 2012; Chiou et al., 2017). We find that Efc25 is recruited to dividing cell ends in a Rax1-dependent manner. However, whether the Rax1/2 complex interacts with other landmarks that play a similar role to Bud8/9 or instead directly binds Efc25 is not yet clear. While the function and basic pathway of the Rax1/2 complex appear to be conserved, many of its interactors identified in *S. cerevisiae* lack homologs in fission yeast. Even Efc25 is not the ortholog of Bud5 but is instead more similar to Cdc25, a different Ras GEF in *S. cerevisiae* (Tratner et al., 1997). These differences suggest that the Rax1/2 complex may have adapted to interact with different proteins while maintaining its role as a memory of growth. Our data with acceptor photobleaching FRET suggest that Rax1 physically interacts with Efc25. However, given the low abundance and the membrane localization of these proteins, traditional co-immunoprecipitation approaches fail to show an interaction. Further biochemical investigations are needed to verify this interaction.

Our results also demonstrate a role for Ras in initiating the site of polarization in fission yeast. Ras-GTPases regulate the site of polarization in multiple fungal species, with the Ras-like GTPase Rsr1 also regulating bud-site selection in *C. albicans*, and *Y. lipolytica* (Hausauer et al., 2005; Li et al., 2014). Rsr1 is also important to maintain the site and direction of polarized growth in fungal hyphae, as loss of Rsr1 leads to random branch initiation and curved hyphae in *C. albicans* and *Ashbya gosspyii* (Hausauer et al., 2005; Bauer et al., 2004). Furthermore, Ras activity has been shown to determine the direction of cell migration in *Dictyostelium* and neutrophils (Sasaki et al., 2004; Zhang et al., 2008; Kortholt et al., 2013; Pal et al., 2023). This suggests that initiation and maintenance of the site of polarization may be a conserved function of Ras GTPases in eukaryotes.

We find that Rax1 regulates Efc25 localization and Ras1 activity primarily in dividing cells, as the level of active Ras1 at interphase cell ends does not change with loss of *rax1*. The timing of Ras1 activation at the old end of dividing cells that we observe is similar to that of the reactivation of the NDR/Orb6 kinase pathway after cytokinesis (Gupta et al., 2014). Previous work has found that Orb6 regulates Ras1 activity by inhibiting translational repression of Efc25 (Nuñez et al., 2016; Chen et al., 2019). Rax1-dependent recruitment of Efc25 may coincide with de-repression of its translation at the end of cell division through increased Orb6 activity, ensuring newly synthesized GEF is targeted to the old end. Our results suggest that rather than the total level of Efc25 at the old end determining its dominance, the direction of its recruitment may be an important factor, as we do not observe any effect on bipolarity from overexpression of Efc25 alone. This may offer a potential explanation for our finding that binding of GBP-mScarlet-Efc25 to Tea1-GFP increases bipolarity, as the dynamics and stability of Efc25 may be altered when bound to Tea1.

Ras1 has previously been shown to promote polarization of Cdc42 through recruitment and activation of Scd1 (Lamas et al., 2020). The Ras1-Cdc42 morphogenesis pathway is also conserved in another fission yeast, as *Schizosaccharomyces japonicus* requires Ras1, Efc25, and the Cdc42 pathway for proper cell shape and hyphal development (Nozaki et al., 2018). We find that in cells lacking Ras1-GTP at dividing cell ends, activation of Cdc42 is delayed and the competition between the two ends is shifted, decreasing the old end’s dominance. With a weaker old end, the new end can activate Cdc42 and initiate growth earlier, leading to the two ends becoming more equal. Interestingly, growth rate is not directly correlated with the amount of Cdc42-GTP at the end, as there was no significant change in Cdc42 activity at the old end despite the lower growth rate in *rax1Δ* cells. This is consistent with other work which found that while increased Cdc42 activity correlated with increased growth, similar levels of active Cdc42 could produce different growth rates and Cdc42 activity varied over the same speed of growth (Taheraly et al., 2020). Our results suggest that the timing of growth along with competition are a significant factor in determining the growth rate of each end, with a delay in onset of growth decreasing the maximal growth rate of that end. This pattern can be seen even in the natural variation of growth among wildtype fission yeast cells, with the old-end growth rate increasing as NETO is delayed (Taheraly et al., 2020). The increased monopolarity in many cytokinetic mutants is also consistent with these findings, as the competitive advantage of the old end increases while new-end growth is inhibited by incomplete division (Bohnert and Gould, 2012).

While *S. pombe* demonstrates a preference for initiating growth at ends with an inherited history of growth, the biological significance for this pattern is unclear. Precocious bipolarity has been associated with decreased cell wall integrity, suggesting early growth at the recently divided end may weaken the cell wall (Koyano et al., 2015). Old-end dominance may also be beneficial under low-nutrient conditions. The timing of NETO is altered in different media, with new-end growth occurring later in minimal media compared to media supplemented with yeast extract (Mitchison and Nurse, 1985). Recent work in our lab has found that stress response pathways prevent precocious bipolarity and that these pathways are downregulated in the transition to bipolar growth (Pathak and Das, 2025). During growth in high cell densities and decreased nitrogen availability, *S. pombe* can form pseudohyphal invasive filaments that are highly monopolar at the ends (Amoah-Buahin et al., 2005; Dodgson et al., 2010). Fission yeast mutants that are more monopolar due to a failure to initiate NETO show more invasive pseudohyphal growth, while cells with premature NETO are less able to form pseudohyphae (Bohnert and Gould, 2012; Bohnert et al., 2020). Initiating growth at the old end may bias growth outwards, away from other cells and potentially towards fresh nutrient sources. Persistence in the direction of migration has been shown to allow chemotaxing cells to search a larger area with higher efficiency (Li et al., 2008), and directed outward growth of filaments may provide similar benefits.

The ability to retain a memory of polarity after it is altered or lost is found across eukaryotic life in many different forms. This memory allows cells to maintain correct polarization through changes in development and the surrounding environment. Even with cell rounding, cortical landmarks can remain asymmetrically localized during division, allowing for positioning of the mitotic spindle, determining cell fate, or re-establishing polarity based on the shape of the mother cell (Théry et al., 2005; Minc and Piel, 2012; Akanuma et al., 2016; Boubakar et al., 2017). Memory is also important in migrating cells, allowing for directed migration towards a chemical signal even during fluctuations or temporary loss of the signal (Skoge et al., 2014; Prentice-Mott et al., 2016; Haastert, 2021). Our work characterizes how fission yeast maintain cell shape across repeated cell divisions by utilizing Rax1/2 as a landmark to reinitiate polarized growth at sites where growth has previously occurred. Our findings also demonstrate how regulation of the timing of Cdc42 activation through memory of growth impacts competition between the two ends.

## Materials and Methods

### Strains and cell culture

*S. pombe* strains used in this study are listed in Table S2. All strains are isogenic to the original strain PN567. Standard methods for genetic manipulation and analysis were used (Moreno et al., 1991). Cells were grown exponentially in rich yeast extract (YE) media at 25°C unless otherwise specified. To express constructs containing the *nmt41* promoter, cells were grown in Edinburgh minimal medium (EMM) with added adenine and uridine (225 µg/ml). Cultures were grown for at least three rounds of eight generations before imaging.

### Strain construction

Deletion of *rax1* was performed as described (Krawchuk and Wahls, 1999). Primers were designed to amplify flanking regions of homology 270 bp immediately upstream of the start codon (forward – 5’-GTATCAAGTAAAGCTTACTA-3’; reverse – 5’- TCAGACAACCACCTCAAGAC-3’) and 250 bp immediately downstream of the stop codon (forward – 5’-CCTGTCGATTTCGTTCCATT-3’; reverse – 5’- CAAAACAGTATAGAATGATT-3’) of *rax1* (SPAC23G3.05c). Coding sequences for all genetic manipulation of *S. pombe* was obtained from PomBase (Wood et al., 2012; Rutherford et al., 2024). The reverse primer for the 5’ flanking region and the forward primer for the 3’ flanking region also contained pFA6a specific sequences as described (Krawchuk and Wahls, 1999). The megaprimers were then used to amplify *kanMX* from the pFA6a-kanMX vector and the product was transformed into wildtype cells. Deletion of *rax1* was confirmed by PCR.

C-terminal tagging of Rax2 with GFP and mScarlet was performed by PCR amplification of 491 bases of the *rax2* coding sequence directly upstream of the stop codon by primers with 5’ overlap homologous to the pFA6a vector (GFP forward – 5’- GCCAGCTGAAGCTTCTGTTGAACTCTGAAGGG-3’; GFP reverse – 5’- TTAATTAACCCGGGGATCCGCTGGGTCTTTAATTCTTTTA-3’; mScarlet forward – 5’- GAACGCGGCCGCCAGTGTTGAACTCTGAAGGG-3’; mScarlet reverse – 5’- CGTTAATTAACCCGGGGCCTCCGCCCCCTCCACCGCCACCCTGGGTCTTTAATTCT TTTA-3’). The amplified fragments were then assembled with either pFA6a-GFP-kanMX digested with BsiWI and SalI or pFA6a-mScarlet-natMX digested with PvuII and BamHI using the NEBuilder HiFi DNA Assembly Cloning Kit (#E5520S, New England Biolabs). The assembled plasmids were then linearized by digestion with SwaI and transformed into wildtype cells.

N-terminal tagging of Rax1 with mNeonGreen was performed using the CRISPR/Cas9 method described in (Fernandez and Berro, 2024). The CRISPR target site and 20bp guide RNA sequence (reverse strand, 5’-AACTTCTGAAACTCTCGGGG-3’) near the start of the *rax1* coding sequence were found using CRISPRdirect (Naito et al., 2015). The donor DNA sequence consisted of regions of homology directly upstream and downstream of the *rax1* coding sequence start codon with point mutations in the PAM and gRNA sequence flanking the coding sequence of mNeonGreen along with a 5-glycine linker. pJB166 (gift from Julien Berro, Addgene #86998) digested with CspCI, the assembled fragment containing the gRNA sequence with flanking regions homologous with pJB166, and the donor DNA were transformed into a *fex1Δfex2Δ* strain (JB355, gift from Julien Berro).

C-terminal tagging of Efc25 with mScarlet was performed by PCR amplification of 630 bases of the *efc25* coding sequence directly upstream of the stop codon by primers with 5’ overlap homologous to the pFA6a vector (forward – 5’- GAACGCGGCCGCCAGTTTAGAAGCTGGGTTAGTCG-3’; reverse – 5’- CGTTAATTAACCCGGGGCCTCCGCCCCCTCCACCGCCACCAGAAGAACGAGGTTC AAGTG-3’). The amplified fragment was then assembled with pFA6a-yomRuby3-natMX (gift from James Moseley) digested with PvuII and AscI to remove the yomRuby3 sequence and the mScarletI coding sequence amplified from pFA6a-mScarletI-kanMX (gift from Julien Berro, Addgene plasmid # 136690) (forward – 5’- CGGATCCCCGGGTTAAT-3’; reverse – 5’-CGCTTATTTAGAAGTGGCG-3’) by NEBuilder HiFi DNA Assembly (#E5520S, New England Biolabs). The assembled plasmid was linearized by MfeI and transformed into wildtype and mNG-Rax1 cells.

To overexpress *efc25*, the vector pJK148-nmt41-mNeonGreen-Efc25-leu1+ was constructed and packaged by VectorBuilder (vector ID: VB240703-1004qpp). The plasmid was linearized with Bsu36I and integrated into the *leu1-32* locus of *efc25Δ* cells. Efc25 was tagged with GBP-mScarlet by replacing the mNeonGreen from pJK148-nmt41-mNeonGreen-Efc25-leu1+ with the GBP sequence amplified from pJK148-Pnmt41-GBP-mCherry(C)-leu1+ (gift from Quanwen Jin, Addgene plasmid # 89069) (forward – 5’-TGTTAAATCAGCCACCATGGCTGATGTCCAACTG-3’; reverse – 5’- TGCTCACCATGCTAGAGGAGACGGTGAC-3’) and mScarletI amplified from pFA6a- mScarletI-kanMX (gift from Julien Berro, Addgene plasmid # 136690) (forward – 5’- TCTAGCATGGTGAGCAAGGGCGAG-3’; reverse – 5’- TCCACCGCCACCCTTGTACAGCTCGTCCATG-3’) using the NEBuilder HiFi DNA Assembly Cloning Kit (#E5520S, New England Biolabs). The assembled plasmid was linearized by Bsu36I and integrated into the *leu1-32* locus of *efc25Δ rax1^+^* cells as well as *rax1Δefc25Δ* cells with and without Tea1-GFP.

### Microscopy

Unless otherwise stated, microscopy was performed at room temperature using a 3i spinning disk confocal using a Zeiss AxioObserver microscope with an integrated Yokogawa spinning disk (Yokogawa CSU-X1 A1 spinning disk scanner) and a 100×/1.49 NA objective. Images were acquired with a Teledyne Photometrics Prime 95b back-illuminated sCMOS camera (Serial No: A20D203014) using SlideBook (3i Intelligent Imaging innovations).

Imaging of calcofluor staining, Rax2-GFP in Supplemental Figure S1B, and timelapse imaging of CRIB-3xGFP, Scd2-GFP, and RasAct^GFP^ was performed using a Nikon Ti2 Eclipse wide-field microscope with a 100×/1.49 NA objective with fluorophores excited using an AURA Light Engine system (Lumencor). Images were acquired with an ORCA-FusionBT digital camera (Hamamatsu Model: C15440-20UP Serial No: 500428) using Nikon NIS Elements (Nikon).

Brightfield imaging of cells for growth analysis was performed using an Olympus IX83 microscope with a VTHawk two-dimensional array laser scanning confocal microscopy system (Visitech International) and a 100×/1.49 NA UAPO lens (Olympus). Images were acquired with a Hamamatsu electron-multiplying charge-coupled device digital camera (Hamamatsu EM-CCD Digital Camera ImageM Model: C9100-13 Serial No: 741262) using MetaMorph (Molecular Devices).

### Calcofluor staining

To visualize cell wall growth, cells were stained in YE liquid containing 50 µg/ml of Calcofluor White (18909, Sigma-Aldrich) at room temperature. Cells were categorized as monopolar or bipolar based on the intensity of staining at the ends and the position of the most recent birth scar.

### Growth analysis

To measure the timing of new-end growth and the growth rate at each end, cells were imaged for 8 hours with an interval of 2 minutes to capture multiple generations. Cells were plated on 35mm glass bottom dishes (MatTek) overlaid with YE containing 0.7% agarose and 100 µM ascorbic acid to prevent phototoxicity. To determine the growth rate, the total length of the cell and the length from the tip of the new end to the birth scar were measured at each time point. NETO was defined as the timepoint when new-end length began consistently increasing. Average growth rate was calculated by dividing the change in length from the onset of NETO to the final length of the cell before division by the amount of time elapsed.

### FRAP imaging and analysis

To inhibit cell growth, cells were treated with YE liquid containing 100 µM CK666 (SML006-5MG, Sigma-Aldrich) dissolved in DMSO (317275-100ML, EMD Millipore) or 100 µM Brefeldin A (B7450, Invitrogen) dissolved in DMSO for 30 minutes prior to imaging. Cells were treated with 1% DMSO as a control. After 30 minutes, cells were plated on 35mm glass bottom dishes (MatTek) overlaid with YE containing 0.7% agarose, 100 µM ascorbic acid, and the appropriate drug treatment or control. Cells were then imaged for 60 minutes at 1 minutes intervals. Rax2-GFP was imaged with 200 ms exposure and 50% laser power at 488 nm wavelength, while mNG-Rax1 was imaged with the same settings except using 70% laser power. After the third timepoint, a 10 x 20 pixel area (1.1 x 2.2 µm) at one of the cell ends was bleached with 20% laser power for 5 ms with 3 repetitions (15 ms total). Acquisition then resumed until the end of the 60-minute timelapse. Full scale normalization of fluorescence recovery data and calculation of the mobile fraction was performed using EasyFRAP-web (Koulouras et al., 2018).

### Fluorescent intensity image acquisition and analysis

Image analysis was performed in Fiji (Schindelin et al., 2012). For still images of fluorescent proteins, z-series of 16 slices at an interval of 0.4 µm were imaged. To measure localization of CRIB-3xGFP, Scd1-mNG, Scd2-GFP, and RasAct^GFP^ at cell ends, images were sum projected and a cell-free region was used for background subtraction. Freehand ROIs were drawn around the cell ends and mean intensity was measured. Tip fraction was calculated as the mean intensity at each end divided by the total end intensity, which was the sum of the mean intensity at both ends of the cell.

To measure Efc25-GFP and mNG-Efc25 intensity, the four middle slices of the z-series were sum projected and background subtraction performed as previously described. Mean intensity was measured using a 4-pixel wide segmented line along the cell cortex from the edge of the division site, across the cell tip, to the opposite side. The centers of the plot profile from each end were then aligned and averaged by binning length into 0.1 µm intervals and calculating the mean and standard deviation for each bin. Intensity was normalized by calculating the mean intensity along the cell side between 3.5 – 5.5 µm from the cell tip for each individual end and subtracting it from the mean intensity, setting negative intensity values to zero. Division site intensity was measured using a 10 pixel (1.1 µm) wide line through the middle of the cell. To calculate whole cell intensity, the full z-series was sum projected, then cells were segmented using the Trainable Weka Segmentation Fiji plugin (Arganda-Carreras et al., 2017) and the mean intensity of each cell was measured.

For timelapse imaging of fluorescent proteins, images were taken of a single z-plane in the middle of the cell. Cells were plated on 35mm glass bottom dishes (MatTek) overlaid with YE containing 0.7% agarose and 100 µM ascorbic acid, then imaged for 1 hour with 1 minute intervals. A cell-free region of the image was used for background subtraction. Freehand ROIs drawn at the cell ends and a polygon ROI surrounding the division site were used to measure mean intensity.

### FRET analysis

FRET efficiency was determined by measuring the increase in donor fluorescence after acceptor photobleaching. Cells were plated on 35mm glass bottom dishes (MatTek) overlaid with YE containing 0.7% agarose and 100 µM ascorbic acid. A single z-plane image was acquired, then photobleaching was performed using 50% laser power at 565 nm wavelength for 3 repetitions of 6 ms each (18 ms total) in freehand ROIs drawn around whole cells. A final image was then acquired after photobleaching. To calculate FRET efficiency, 4-pixel wide segmented line ROIs were drawn along the cell cortex at regions of colocalization pre-bleach and mean intensity was measured in the 488 channel in both images acquired pre- and post-acceptor photobleaching. Unbleached cells in the same image were also measured as controls. Post-acceptor photobleaching intensities were corrected for photobleaching of the donor fluorophore. The percent FRET efficiency was calculated by dividing the initial donor fluorescence intensity by the post-bleach intensity, subtracting the result from 1 and multiplying by 100.

### Statistical tests

Statistical significance was calculated using GraphPad Prism. Unless otherwise specified, a two-tailed student’s t-test with Welch’s correction was used when comparing two conditions. A one-way ANOVA (unequal variance) corrected for multiple comparisons was used when comparing three or more conditions. Graphs were made in GraphPad Prism.

## Acknowledgements

We thank Fulvia Verde, Sophie Martin, and Julien Berro for providing strains, and James Moseley, Quanwen Jin, and Julien Berro for providing plasmids. We also thank Bret Judson and the Boston College Imaging Core for infrastructure and support.

This work was supported through an NIH/NIGMS R01GM136847 award to M. Das.

