## Supplementary Material for "Rax1/2 promote memory of growth by early Ras1 activation to maintain polarity across generations"

**Table S1. Percentage of bipolar cells in *rax1*$^{+}$ and *rax1*Δ strains.**

| | % Bipolarity | |
|---|---|---|
| Background | *rax1*$^{+}$ | *rax1*Δ |
| *wt* | 71.9 ± 3.06 | 84.2 ± 2.88 |
| *gef1*Δ | 54.5 ± 5.61 | 79.0 ± 1.52 |
| *scd2*Δ | 22.7 ± 1.89 | 42.0 ± 1.68 |
| *rga3*Δ | 77.6 ± 0.910 | 84.5 ± 0.985 |
| *rga4*Δ | 73.5 ± 1.29 | 85.4 ± 3.53 |
| *rdi1*Δ | 57.8 ± 8.41 | 81.5 ± 6.97 |
| *bud6*Δ | 57.2 ± 6.40 | 67.3 ± 2.16 |

The mean and standard deviation of the percentage of bipolar cells as determined by calcofluor staining. Means calculated from 3 independent experiments (n ≥ 97 cells, N = 3). Values for the wildtype *rax1*$^{+}$ and *rax1*Δ strains are the mean and standard deviation of the plot in Figure 1B.

**Table S2. List of strains used in this study.**

| Strain | Genotype | Origin | Used |
|---|---|---|---|
| PN567 | *h+ ade6-704 leu1-32 ura4-D18* | P. Nurse | Fig 1 A-G,<br>Fig 5 A,B, &<br>Table S1 |
| YMD1866 | *h- rax1Δ::kanMX6 ade6-704 leu1-32 ura4-D18* | This study | Fig 1 C-G |
| YMD1882 | *h+ rax1Δ::kanMX6 ade6-704 leu1-32 ura4-D18* | This study | Fig 1 A,B,<br>Fig 5 A,B, &<br>Table S1 |
| FV0001 | *h+ orb2-34 ade6-M216 leu1-32* | F. Verde | Fig 1 H,I |
| YMD2346 | *rax1Δ::kanMX6 orb2-34 ade6- leu1-32* | This study | Fig 1 H,I |
| YMD2351 | *h- rax2-GFP:kanMX6 ura4-D18 ade6-704 leu1-32* | This study | Fig 2 A-C,F &<br>Fig S1 A,C |
| YMD2246 | *h+ rax2-GFP:kanMX6 ura4-D18 ade6-704 leu1-32* | This study | Fig S1 B |
| YMD2285 | *h- rax1Δ::kanMX6 rax2-GFP:kanMX6 ade6-704 leu1-32 ura4-D18* | This study | Fig S1 B |
| YMD2732 | *h- mNeonGreen-gly8-rax1 fex1+ fex2+ his3+ ura4-D18 ade6- leu1-32* | This study | Fig 2 A,D,E,G<br>& Fig S1 A,D |
| FV1335 | *h+ CRIB-3xGFP:ura4+ ura4-D18 leu1-32* | F. Verde | Fig 3 A,B,E-G,<br>Fig 5 C-E, &<br>Fig S2 A,D |
| YMD1913 | *CRIB-3xGFP:ura4+ rax1Δ::kanMX6 ura4-d18 leu1-32* | This study | Fig 3 A,B,E-G,<br>Fig 5 C-E, &<br>Fig S2 A,D |
| YMD1176 | *h+ scd1-mNG:kanMX6 ade6-704 leu1-32 ura4-D18* | Lab strain | Fig 3 A,C,<br>Fig S2 B,E, &<br>Fig S3 A-C |
| YMD1909 | *h+ rax1Δ::kanMX6 scd1-mNG:kanMX6 ade6-704 leu1-32 ura4-D18* | This study | Fig 3 A,C,<br>Fig S2 B,E, &<br>Fig S3 A-C |
| FV1118 | *h+ scd2-GFP:kanMX6 leu1-32 ura4-D18* | F. Verde | Fig 3 A,D,<br>Fig S2 C,G, &<br>Fig S3 D-F |
| YMD1911 | *h- rax1Δ::kanMX6 scd2-GFP:kanMX6 leu1-32 ura4-D18* | This study | Fig 3 A,D,<br>Fig S2 C,G, &<br>Fig S3 D-F |
| YSM3098 | *h- ura4-D18 leu1-32:RasAct-3GFP:leu1+* | S. Martin (Merlini et al., 2018) | Fig 4 A-G |

| | | | |
|---|---|---|---|
| YMD2397 | *rax1Δ::kanMX6 leu1-32::RasAct-3xGFP:leu1+ ura4-D18* | This study | Fig 4 A-G |
| YMD2621 | *tea1Δ::ura4+ leu1-32::RasAct-3xGFP:leu1+ ura4-D18* | This study | Fig 4 H |
| YMD2525 | *h- efc25Δ::kanMX6 rax1Δ::kanMX6 ura4-D18 leu1+* | This study | Fig 5 A,B |
| YMD2546 | *h+ efc25Δ::kanMX6 ade6-704 leu1-32 ura4-D18* | This study | Fig 5 A,B |
| YMD1268 | *scd2Δ::kanMX6 CRIB-3xGFP:ura4+ ura4-* | Lab strain | Fig 5 C-E |
| YMD2613 | *efc25Δ::kanMX6 CRIB-3xGFP:ura4+ ura4-* | This study | Fig 5 C-E |
| YMD2679 | *rax1Δ::kanMX6 efc25-gfp:kanMX6 ura4-D18* | This study | Fig 6 A-C,G |
| YMD2680 | *efc25-gfp:kanMX6 ura4-D18* | This study | Fig 6 A-C,G |
| YMD2640 | *h+ leu1-32::pjk148-Pnmt41-mNeonGreen-gly8-efc25:leu1+ efc25Δ::kanMX6 ade6-704 ura4-D18* | This study | Fig 6 D-F,H,I |
| YMD2664 | *leu1-32::pjk148-Pnmt41-mNeonGreen-gly8-efc25:leu1+ efc25Δ::kanMX6 rax1Δ::kanMX6 ade6-704 ura4-D18* | This study | Fig 6 D-F,H,I |
| YMD1547 | *h+ leu1-32::pjk148-Pnmt41-leu1+* | Lab strain | Fig 6 I |
| YMD2856 | *h- rax1Δ::kanMX6 leu1-32::pjk148-Pnmt41-leu1+* | This study | Fig 6 I |
| YMD2786 | *h+ efc25Δ::kanMX6 leu1-32::pjk148-Pnmt41-GBP-mScarlet-gly8-efc25-leu1+ ade6-704 ura4-D18* | This study | Fig 7 A,B |
| YMD2788 | *h+ efc25Δ::kanMX6 rax1Δ::kanMX6 leu1-32::pjk148-Pnmt41-GBP-mScarlet-gly8-efc25-leu1+ ura4-D18 ade6-* | This study | Fig 7 A,B |
| YMD2790 | *rax1Δ::kanMX6 efc25Δ::kanMX6 tea1-gfp:kanMX6 leu1-32::pjk148-Pnmt41-GBP-mScarlet-gly8-efc25-leu1+ ade6- ura4-* | This study | Fig 7 A,B |
| YMD2820 | *efc25Δ::kanMX6 tea1-gfp:kanMX6 leu1-32::pjk148-Pnmt41-GBP-mScarlet-gly8-efc25-leu1+ ade6- ura4-* | This study | Fig 7 A,B |
| YMD2847 | *mNeonGreen-gly8-rax1 rax2-gly8-mScarlet:natMX6 fex1+ fex2+ his3+ ura4-D18 ade6- leu1-32* | This study | Fig 7 C,D |
| YMD2733 | *mNeonGreen-gly8-rax1 efc25-mScarletI:natMX6 fex1+ fex2+ his3+ ura4-D18 ade6- leu1-32* | This study | Fig 7 E,F |

| | | | |
|---|---|---|---|
| YMD2739 | *efc25-gly8-mScarletI:natMX6 rax2-gfp:kanMX6 ura4-D18 leu1-32 ade6-* | This study | Fig 7 E,G |
| YMD776 | *h- rdi1Δ::kanMX6 ura4-D18* | Lab strain | Table S1 |
| YMD1894 | *rax1Δ::kanMX6 rdi1Δ::kanMX6* | This study | Table S1 |
| YMD476 | *h- gef1Δ::ura4+ ura4-* | Lab strain | Table S1 |
| YMD1887 | *gef1Δ::ura4+ rax1Δ::kanMX6 ura4-* | This study | Table S1 |
| YSM3131 | *h- rga3Δ::kanMX6 ade6- leu1-32 ura4+* | S. Martin (Gallo Castro & Martin, 2018) | Table S1 |
| YMD1929 | *rax1Δ::kanMX6 rga3Δ::kanMX6 ade6- leu1-32 ura4+* | This study | Table S1 |
| FV244 | *h90 ura4-D18 scd2Δ::ura4+* | F. Verde | Table S1 |
| YMD2443 | *rax1Δ::kanMX6 scd2Δ::ura4+ ura4-D18 leu1- ade6-* | This study | Table S1 |
| FV862 | *h- bud6Δ::kanMX* | F. Verde | Table S1 |
| YMD2619 | *rax1Δ::kanMX6 bud6Δ::kanMX6* | This study | Table S1 |
| YMD664 | *h+ rga4Δ::ura4+ ura4-* | Lab strain | Table S1 |
| YMD2868 | *rax1Δ::kanMX6 rga4Δ::ura4+ ura4-* | This study | Table S1 |
| YLM153 | *h- efc25-gfp:kanMX6 ura4+ leu1+ ade6+* | S. Martin | Strain construction |
| FV2895 | *h- efc25Δ::kanMX6 ura4+ leu1+* | F. Verde (Chen et al., 2019) | Strain construction |
| JB355 | *h- fex1Δ fex2Δ his3-D1 ura4-D18 leu1-32 ade6-M216* | J. Berro (Fernandez & Berro, 2024) | Strain construction |
| FV467 | *h+ tea1Δ::ura4+ ura4-D18 ade6-M210* | F. Verde | Strain construction |
| FV551 | *h- tea1-GFP:kanMX6 ura4- ade6- leu1-* | F. Verde | Strain construction |

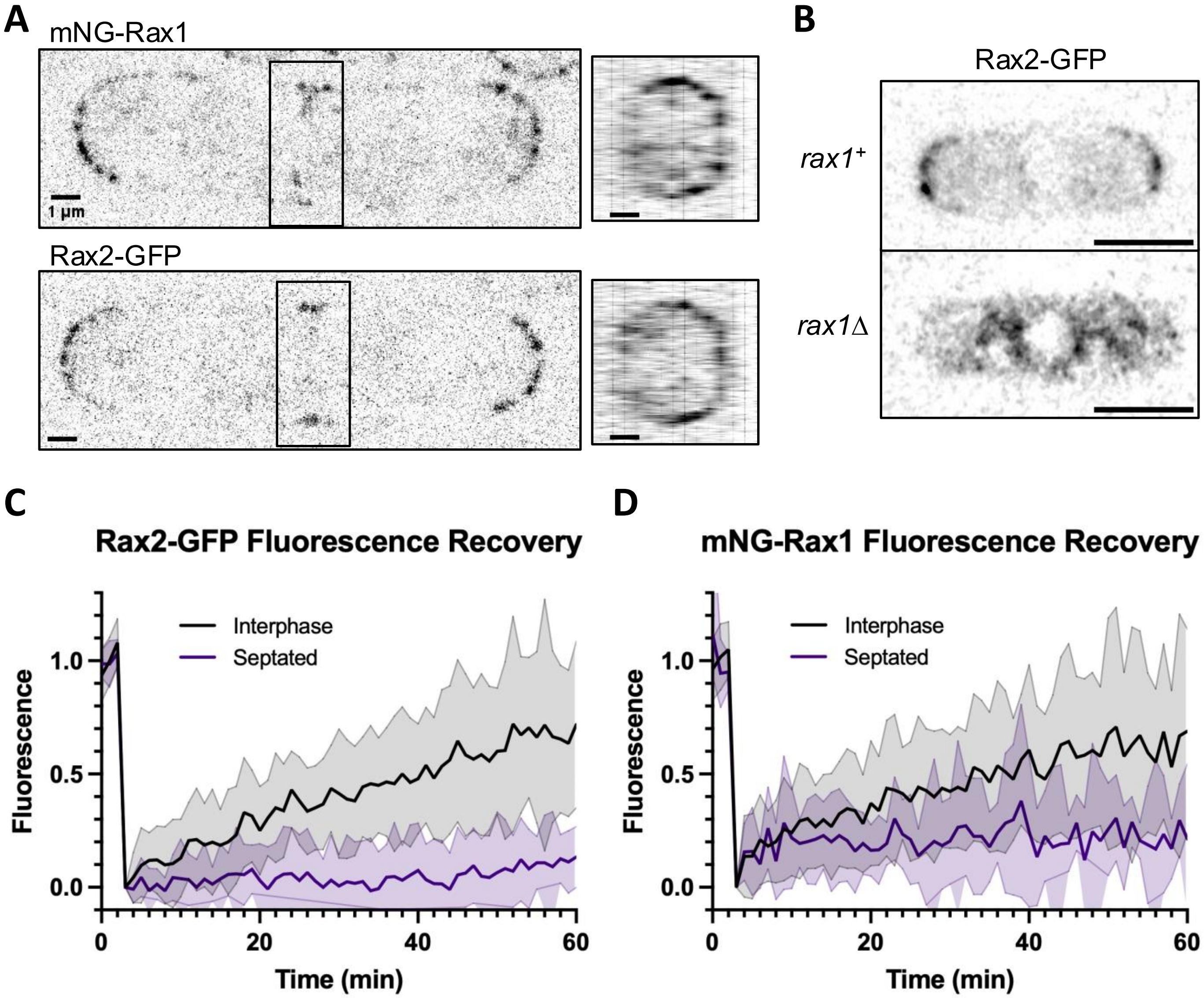


**Figure S1. Localization and fluorescence recovery of Rax1 and Rax2 in dividing cells. (A)** SoRa images of mNG-Ras1 and Rax2-GFP in dividing cells, with a 3D projection of the inset rotated 90º. 2.8x magnification. Scale bars are 1 µm. **(B)** Localization of Rax2-GFP in *rax1*$^{+}$ and *rax1*Δ cells. Scale bars are 5 µm. **(C,D)** Mean and s.d. of normalized fluorescence recovery over time of Rax2-GFP (C) and mNG-Rax1 (D) in DMSO treated cells. (n ≥ 6 cells). Data from interphase cells is the same as the DMSO treated control in Figure 2 F,G.

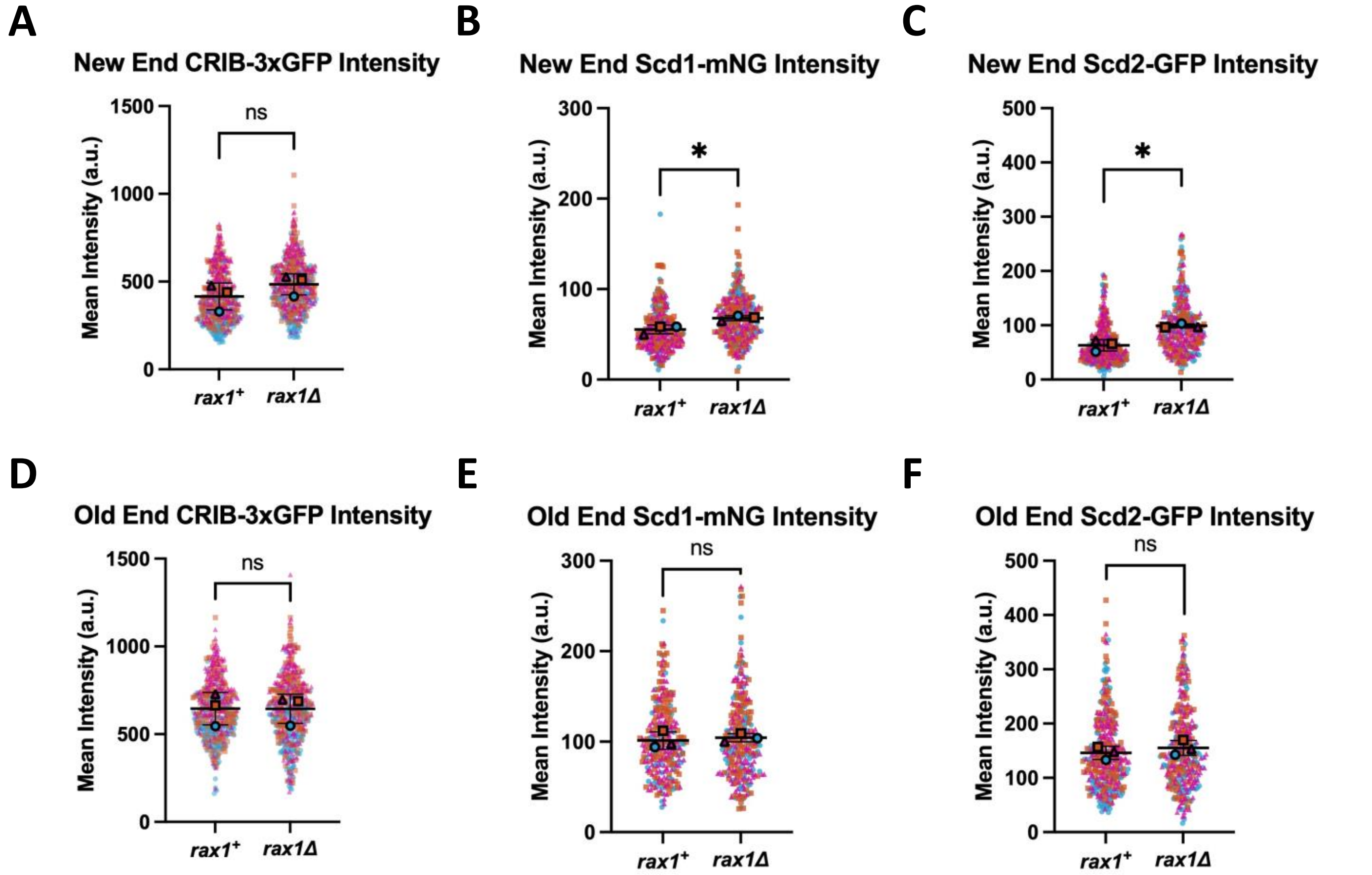


**Figure S2. Loss of *rax1* increases activation of Cdc42 at the new ends.** Intensity of CRIB-3xGFP (A,D), Scd1-mNG (B,E), and Scd2-GFP (C,F) at the new ends (A-C) or old ends (D-F) of *rax1*$^+$ and *rax1*Δ cells. Outlined points represent the mean from one independent experiment, with small points corresponding to individual cells ($n \geq 54$ cells, $N = 3$). Error bars are s.d. Significance was determined using an unpaired, two-tailed t-test with Welch's correction. * = $p < 0.05$, ns = not significant.

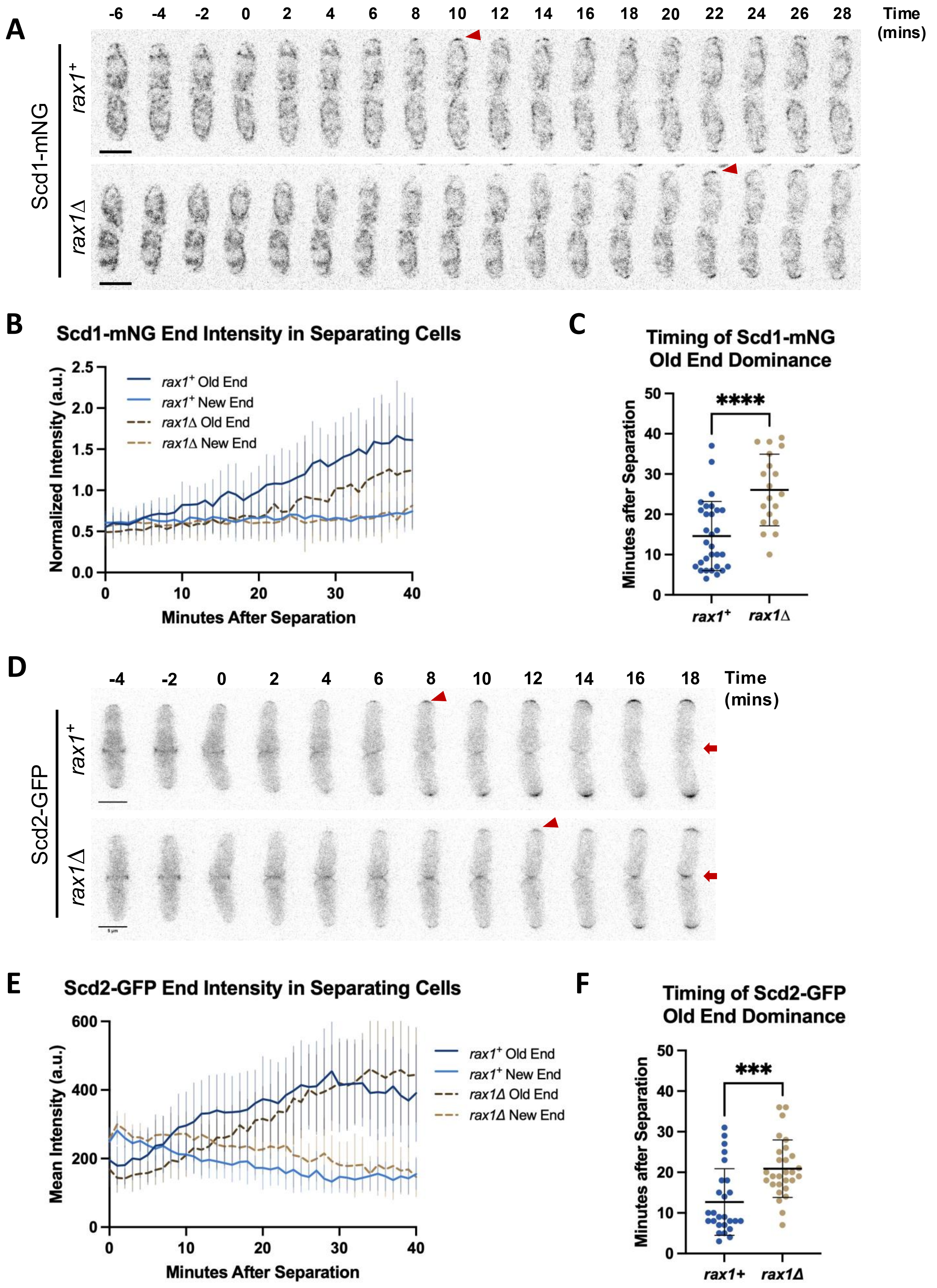

A
-6 -4 -2 0 2 4 6 8 10 12 14 16 18 20 22 24 26 28
Time (mins)
Scd1-mNG
rax1+
rax1Δ
B
Scd1-mNG End Intensity in Separating Cells
Normalized Intensity (a.u.)
Minutes After Separation
rax1+ Old End
rax1+ New End
rax1Δ Old End
rax1Δ New End
C
Timing of Scd1-mNG Old End Dominance
Minutes after Separation
****
rax1+
rax1Δ
D
-4 -2 0 2 4 6 8 10 12 14 16 18
Time (mins)
Scd2-GFP
rax1+
rax1Δ
E
Scd2-GFP End Intensity in Separating Cells
Mean Intensity (a.u.)
Minutes After Separation
rax1+ Old End
rax1+ New End
rax1Δ Old End
rax1Δ New End
F
Timing of Scd2-GFP Old End Dominance
Minutes after Separation
***
rax1+
rax1Δ

**Figure S3. Recruitment of Scd1-mNG and Scd2-GFP to the old ends after cell separation is delayed in *rax1*Δ cells. (A)** Montage of Scd1-mNG in separating *rax1*$^{+}$ and *rax1*Δ cells. Arrowheads mark localization of Scd1-mNG at old ends. Time '0' is onset of cell separation. **(B)** Quantification of mean and s.d. of normalized Scd1-mNG intensity over time at the old and new ends of separating *rax1*$^{+}$ and *rax1*Δ cells ($n \geq 20$). Intensity was normalized by cytoplasmic intensity. **(C)** Quantification of the timepoint at which old-end Scd1-mNG intensity becomes higher than new-end intensity. Data points correspond to individual cells. **(D)** Kymographs of Scd2-GFP in separating *rax1*$^{+}$ and *rax1*Δ cells. Arrowheads mark localization of Scd2-GFP at old ends, while arrows mark the new ends. Time is in minutes from cell separation. **(E)** Quantification of mean and s.d. of Scd2-GFP intensity over time at the old and new ends of separating *rax1*$^{+}$ and *rax1*Δ cells ($n \geq 18$). **(F)** Quantification of the timepoint at which old-end Scd2-GFP intensity becomes higher than new-end intensity. Data points correspond to individual cells. Scale bars are 5 µm. Error bars are s.d. Significance was determined using an unpaired, two-tailed t-test with Welch's correction. *** = $p < 0.001$, **** = $p < 0.0001$.
